# *Odoribacter splanchnicus* mitigates pathogen*-*induced gut inflammation and its associated pathogenesis via its secreted bacteriocin

**DOI:** 10.1101/2024.08.23.609322

**Authors:** Dipasree Hajra, Debapriya Mukherjee, Rhea Vij, Raju S Rajmani, Venkatareddy Dadireddy, Deepakash Das, Tanweer Hussain, Mahipal Ganji, Utpal Tatu, Dipshikha Chakravortty

## Abstract

Foodborne pathogens pose a major global health hazard, worsened by drug-resistant strains. Therefore, designing novel therapeutic strategies is crucial. Here, we identified the protective role of gut-commensal *Odoribacter splanchnicus* (OS) against *Salmonella* in mice by promoting colonization resistance, preserving gut barrier integrity, and preventing acute inflammation and biofilm formation. *In vitro,* both OS and their culture supernatant inhibited *Salmonella* biofilm formation, intracellular proliferation in human intestinal cells, and virulence gene expression. Further, our results depicted that OS’s protective role acts over a broad spectrum as it confers protection against Gram-positive, *Listeria monocytogenes* and Gram-negative, *Salmonella* Typhimurium. Notably, OS conferred protection even when administered post-infection in mice, highlighting its therapeutic potential. Using several biochemical and proteomics approaches, we characterized key OS-secreted molecules that limit intracellular *Salmonella* and *Listeria* replication in human intestinal epithelial cells by regulating key virulence effectors and flagella. Collectively, our study establishes OS as the broad-spectrum protective agent against *Salmonella* and *Listeria* infections, with promising therapeutic potential.

**Author Summary:** Foodborne pathogens such as *Salmonella* and *Listeria* continue to threaten global health, especially with the rise of antibiotic resistance. In this study, we identify the gut commensal bacterium *Odoribacter splanchnicus* (OS) as a potent protectant against these pathogens. Using a mouse model, we show that OS enhances colonization resistance, maintains gut barrier integrity, and prevents both inflammation and biofilm formation during infection. Remarkably, OS also limits pathogen proliferation and virulence gene expression in human intestinal cells. These protective effects are mediated by both live OS and its secreted molecules within culture supernatant. Importantly, OS administration post-infection also conferred protection, indicating strong therapeutic potential. Biochemical and proteomic analyses revealed OS-secreted factors that suppress key virulence and motility pathways in pathogens. Our findings establish *Odoribacter splanchnicus* as a promising broad-spectrum biotherapeutic agent against major foodborne infections.

**GRAPHICAL ABSTRACT:** 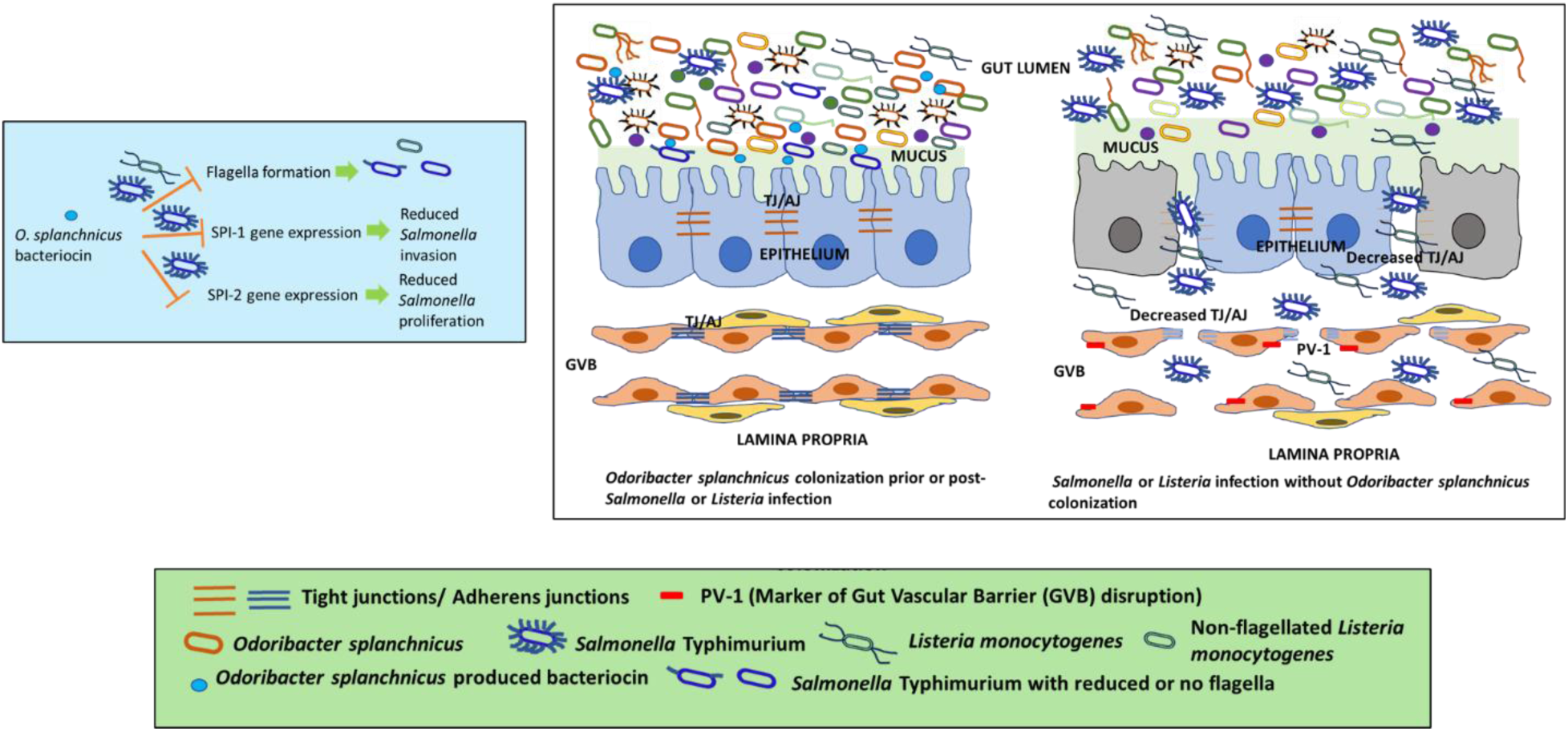

## INTRODUCTION

The existence of the gut vascular barrier (GVB) acts as a protective barrier against the entry of microorganisms into systemic blood circulation (1, 2). Enteric pathogens such as *Salmonella enterica* can cross the mucus and intestinal epithelial barrier, ultimately leading to GVB damage and systemic dissemination (1). The gut microbiota aids in the maintenance of the intestinal epithelial barrier by the production of several metabolites such as short-chain fatty acids (SCFAs), indoles, and polyphenol metabolites that contribute to epithelial barrier integrity and turnover by regulating the expression of tight-junction (TJ) genes (3). Additionally, these SCFAs have the potential to exert anti-inflammatory properties by regulating cytokine production, mucosal immune cell migration, and secretion of antimicrobial peptides (AMPs) by intestinal epithelial cells and macrophages (4–6).

*Salmonella* enters the body through the ingestion of contaminated food and water, resisting the acidic stomach pH with acid tolerance response (ATR) genes, and finally reaches the lumen of the small intestine(7). The pathogen gains entry into the non-phagocytic epithelial cells via *Salmonella* Pathogenicity Island-1 (SPI-1) effectors to breach the intestinal barrier (8). Once traversing through the mucosal lining of the gut epithelium, it reaches the lamina propria (8). In the lamina propria, *Salmonella* encounters various host immune cells like neutrophils, macrophages, and dendritic cells. Macrophages, dendritic cells, and neutrophils are responsible for successful dissemination throughout the body through the reticulo-endothelial system (RES) to the secondary organs such as Mesenteric Lymph Node (MLN), liver, and spleen(8). *Salmonella* relies on the SPI-2 secretion for its intracellular and intravacuolar life by injecting various virulence factors into the host cell cytosol (9, 10).

*Odoribacter splanchnicus* (OS, belonging to the order Bacteroidales, is a common member of the human intestinal microbiota. *Bacteroidetes* is one of the major commensal phyla dominating the healthy mouse as well as the human gut (11). It is a Gram-negative, anaerobic, non-motile and non-spore-forming bacterium. A decrease in *Odoribacter* abundance has been linked to different microbiota-associated diseases, such as non-alcoholic fatty liver disease, cystic fibrosis, and inflammatory bowel disease (IBD)(12). A recent report indicated the ability of OS and its effectors transported in Outer membrane vesicles (OMVs) to exhibit anti-inflammatory effects on the gut epithelium(12). Further, OS has been implicated in conferring resistance against colitis and colorectal cancer(13). This led us to hypothesize that *O. splanchnicus* might ameliorate the inflammatory disease conditions associated with *Salmonella* infection and dissemination. Here, in the present study, we aim to understand whether colonization with the gut commensal OS, could ameliorate *Salmonella-*induced inflammation and pathogenic dissemination.

Our study shows that OS colonization promotes colonization resistance of *Salmonella*, and ameliorates systemic dissemination. OS ameliorates acute *Salmonella*-induced inflammation and its associated pathogenesis by promoting colonization resistance, resulting in decreased *Salmonella* invasion of gut tissue, the preservation of tight junctions, and GVB maintenance. Further, we demonstrate that not only live OS but its culture supernatant inhibits *Salmonella in vitro* biofilm formation, intracellular proliferation in human intestinal epithelial cells, and virulence gene expression. Subsequently, our results depict that the specific protective role of OS acts over a broad spectrum as it confers protection against both flagellated Gram-positive, *Listeria monocytogenes* and Gram-negative, *Salmonella* Typhimurium. Furthermore, OS conferred protection even after its administration to mice post-establishment of infection highlighting its therapeutic potential. Subsequently, we identified the key OS-secreted bacteriocin protein responsible for limiting intracellular *Salmonella* and *Listeria* replication in human intestinal cells by regulating virulence genes such as flagella. Overall, our study implicates the broad-spectrum protective role of OS in mitigating foodborne infections caused by *Salmonella* and *Listeria* pathogenesis.

## MATERIALS AND METHODS

### Bacterial Strains and Culture Conditions

*Salmonella enterica* serovar Typhimurium (STM) strain ATCC 14028S, STM ATCC 14028S constitutively expressing mCherry (pFPV25.1-mCherry-Amp^R^, RFP), *Salmonella enterica* serovar Typhi CT18 (STY), laboratory stock of *Enterococcus faecalis* strain API 142 MTCC 439 (ATCC 35550) (Ef), *Listeria monocytogenes* MTCC 657 (LM) expressing mCherry (pFPV25.1-mCherry-Amp^R^), and *Odoribacter splanchnicus* (OS) strain NCTC 10825 [1651/6] from ATCC were used in this study. 4% paraformaldehyde-fixed-OS was used as killed bacteria control. The STM, LM, and Ef bacterial strain was grown overnight in Miller LB broth (Himedia, M1245) at 37 ^0^C under 160 rpm shaking conditions. OS was cultured under anaerobic conditions (anaerobic chamber) in tryptic soy broth (Himedia, M-011) at 37°C for 48-72h after its revival from glycerol stock.

### Preparation of Cell-free Bacterial Culture Supernatant

The overnight grown OS culture in tryptic soy broth was centrifuged at 10,000rpm for 10 min and the supernatant was harvested. The collected supernatant was further subjected to another two rounds to centrifugation to remove any residual cells. The supernatant was subsequently filtered using a 0.2 µm filter (Cole-Parmer PTFE Syringe Filters) and stored at − 20°C.

### Animal Experiments

The animal experiments were conducted in accordance with the approved guidelines of the institutional animal ethics committee at the Indian Institute of Science, Bangalore, India (Registration No: 48/GO/ReRcBiBt-S/Re-L/99/CPCSEA). All procedures involving the use of animals were performed according to the Institutional Animal Ethics Committee (IAEC)-approved protocol (CAF/Ethics/853/2021). 6–8 weeks old adult C57BL/6 male mice were used in this study (bred, housed, and provided (purchased) by Central Animal Facility, Indian Institute of Science, Bangalore). Male mice were used to minimize variability associated with estrous cycle–dependent hormonal fluctuations that can modulate innate and adaptive immune responses and influence infection outcomes. Using a single sex reduced experimental variability and improved statistical power to detect infection-induced phenotypes within the scope of this study. After one day of OS colonization (10^5^ CFU/mice via oral gavage), the mice were infected with 10^6^ (organ burden) CFU or 10^8^ CFU (for assessing survival) of *S*. Typhimurium, and 10^8^ (organ burden) or 10^9^ (survival) CFU of *L. monocytogenes* via oral gavage. The colonization of *O. splanchnicus* was confirmed via RT-qPCR mediated quantitation of OS 16S rRNA from the mice fecal (3 fecal pellets harvested from the intestine and resuspended in Trizol) and ileal samples at 6^th^ day post-OS colonization. For post-infection OS administration studies, OS was orally administered (10^5^ CFU/ mice) post 1 or 3dpi. 5-day post-infection, mice were euthanized, dissected out, and blood and organs were harvested. The organs were homogenized using a bead-beater. The bacterial organ load was enumerated by plating the tissue homogenates on *Salmonella-Shigella* (SS) agar plates for *Salmonella* or LB plates containing 50µg/ml of Ampicillin for LM. The harvested tissues and the fecal samples were collected in protein lysis buffer (RIPA) and TRIzol (Takara) for protein and RNA extraction respectively and were stored at −80°C. For calculating percentage survival, the infected mice cohorts were monitored till fatality and fatalities were reported.

### Blood collection and serum preparation

Blood collection was performed on the 5^th^ day post-infection via cardiac puncture post-euthanization of the mouse. The collected blood was allowed to clot by incubating the vials at room temperature for 45 minutes. After allowing the standing time of 45 mins, centrifugation was performed at 2000g for 15 minutes in a refrigerated centrifuge. Following the centrifugation, the clear, yellowish supernatant was transferred to an autoclaved microfuge tube and stored at −80°C till further downstream processing.

### ELISA

Cytokines in mice serum were quantified according to the manufacturer’s instructions(10, 14). Briefly, 96-well ELISA plates of IL-6 ELISA kit (BD Bioscience- Cat No-555240) were coated overnight with capture antibody at 4°C. Following day, plates were washed with 0.1% Tween-20 containing PBS and blocked with 10% FBS for 1 h. Following blocking, wells were washed and incubated with 50µl of serum samples in duplicates for 2 h at room temperature. Subsequently, plates were washed and incubated with detection antibody and enzyme reagent for 1 h at room temperature (BD Bioscience). TMB (Sigma) was used as a substrate and reactions were stopped with 2 N H_2_SO_4_.

Absorbance was measured at 450 nm wavelength in Tecan Plate reader and the concentration of cytokines were interpolated from a standard curve.

### Quantitative Real Time PCR

Total RNA was isolated from the ileal section of the intestinal tissues or feces harvested in TRIzol (Takara) as per manufacturer’s protocol. RNA was quantified using Nano Drop (Thermo-Fischer scientific).3 µg of total isolated RNA was subjected to DNaseI (Thermo Fischer Scientific) treatment at 37°C for 1 hr followed by heat inactivation at 75°C in the presence of 5mM concentration of EDTA for 10 mins. The mRNA was reverse transcribed to cDNA using oligo (dT)_18_ primer, random hexamer primer, buffer, dNTPs and reverse transcriptase (Takara) as per manufacturer’s protocol. The expression profile of target gene was evaluated using specific primers (Table-S2) by using SYBR green RT-PCR master mix (Takara) in Applied Biosystems (Quantstudio 5) or BioRad Real time PCR instrument. β-actin was used as an internal control for mammalian genes and for bacterial genes 16S rRNA was used. All the reaction was setup in 384 well plate with three replicates for each sample.

### Western blotting

The harvested tissue samples were lysed in ice-cold RIPA buffer (50 mM Tris-Cl, pH 8.0, 150 mM NaCl, 5mM EDTA, 1% NP-40, 1 % Sodium deoyxycholate, 1% SDS) containing protease inhibitor cocktail (Roche). Total protein was estimated using Bradford reagent (BioRad). Protein samples were denatured in **Laemmli SDS loading buffer** at 95°C for 10 min and then subjected to separation on 12% SDS-PAGE gels followed by transfer onto activated 0.45µm PVDF membrane. Finally, the membrane was blocked with 5% skimmed milk in TBST (Tris Buffered Saline containing 0.1% Tween-20) and probed with appropriate primary (Table-S3) and secondary HRP-conjugated antibodies. The blots were developed using ECL substrate (Bio-Rad) using a Chemidoc (BioRad). All densitometric analysis was performed using ImageJ software.

### Immunohistochemistry

The harvested tissues were fixed in 4% paraformaldehyde, embedded in paraffin and 5 μm thick paraffin sections were collected on poly-lysine coated glass slides. The sectioned tissue samples were deparaffinized using standard protocol. Briefly, the slides containing the paraffinized tissues were immersed in xylene for 10 minutes and this step was repeated twice. After the xylene treatment, the tissues were subjected to absolute alcohol treatment for 5 minutes twice, followed by 95% and 70% alcohol treatment for 5 minutes. The slides were rinsed in water followed by boiling step in water for 2 minutes and then dried. For in vivo estimation of biofilm, deparaffinized tissue samples were stained with Calcofluor White (Sigma) stain for 20 min at RT. All immunofluorescence images were obtained using Zeiss multi-photon 880 microscope and were analyzed using ZEN black 2012 software.

### Haematoxylin and Eosin Staining

The harvested ileal tissues were collected and fixed using 3.5% paraformaldehyde. The fixed liver was then dehydrated using a gradually increasing concentration of ethanol and embedded in paraffin. 5μm sections were collected on coated plates. Sections were further rehydrated and then stained with hematoxylin and eosin. Images were collected in a Leica microscope.

The scoring of the pathological changes is graded as 0-4; for each of the pathological lesions (Severe/ marked −4; Moderate −3; Mild −2; Minor/minimum −1; No pathology −0) considering the intestinal inflammation and associated villi injury as described previously (15, 16).

### Cell Culture

Human colorectal adenocarcinoma cell line Caco-2 were maintained in Dulbecco’s Modified Eagle’s Media (DMEM) (Lonza) and supplemented with 10% FCS (Fetal calf serum, Gibco) and 1% non-essential amino acid solution (Sigma Aldrich) at 37°C temperature in the presence of 5% CO_2_ in humidified incubator. Prior to each experiment, cells were seeded into 24 well or 6 well plate as per requirement at a confluency of 60%.

### Intracellular proliferation or gentamicin protection assay

Caco-2 epithelial were infected with 3 to 4 h old log phase STM or LM or STY bacterial culture, STM (or LM or STY) with OS, STM (or LM or STY) in presence of *O. splanchnicus* supernatant and STM along with PFA-fixed OS at an MOI of 10. For attachment of bacteria to the cells, tissue culture plates were subjected to centrifugation at 600xg for 5 min and incubated at 37 ^0^C humified incubator with 5% CO_2_ for 25 min. Following infection of the cells with STM at an MOI of 10, cells were treated with DMEM (Sigma) + 10% FBS (Gibco) containing 100 μg/ml gentamicin for 1 hr. Subsequently, the gentamicin concentration was reduced to 25 μg/ml and maintained until the specified time point. Post 2hr and 16hr post-infection, cells were lysed in 0.1% triton-X-100. Lysed cells were serially diluted and plated on *Salmonella-Shigella* (SS) or LB agar to obtain colony-forming units (CFU). Percent invasion was determined using the following formula (17, 18):

**Percent invasion= [CFU at 2 h]/ [CFU of pre-inoculum] X 100**

Fold proliferation was calculated as CFU at 16hr divided by CFU at 2hr.

**Fold Proliferation= [CFU at 16h] / [CFU at 2h]**

### Immunofluorescence confocal microscopic studies

At the specified time points post infection with mCherry-tagged STMAS, cells were fixed with 3.5% paraformaldehyde for 15 min. Primary antibody staining was performed with specific primary antibody in the presence of a permeabilizing agent, 0.01% saponin (Sigma) dissolved in 2.5% BSA containing PBS at 4°C for overnight or for 6hr at room temperature (RT). Following this, cells were washed with PBS stained with appropriate secondary antibody tagged with fluorochrome for 1 hr at RT. This was followed by DAPI staining and mounting of the coverslip onto a clean glass slide using the mounting media containing the anti-fade agent. The coverslip sides were sealed with a transparent nail paint. All immunofluorescence images were obtained using Zeiss Multiphoton 880 and were analyzed using ZEN Black 2012 software.

### Growth Curve analysis

For *in vitro* growth curve experiments a single colony of STM and OS was inoculated in 5mL of LB broth singly or in co-culture and grown overnight at 37°C in anaerobic jar. In another set, STM was inoculated in 5mL of LB in presence of OS supernatant. Overnight-grown stationary phase bacteria were sub-cultured at a ratio of 1: 100 in freshly prepared LB in presence or absence of OS supernatant and kept at 37°C. At different time intervals, aliquots were taken for measurement of [OD]600nm by TECAN 96 well microplate reader and were plated for CFU enumeration.

### Biofilm-inducing growth media and conditions

LB medium lacking NaCl and containing 1% casein enzyme hydrolysate (Hi-Media, India) and 0.5% yeast extract (Hi-Media, India)) was used as biofilm media as per previous literature (19). Overnight grown culture was subcultured in 2 ml biofilm medium at the dilution of 1:100 in a flat-bottom 24-well plate (Tarsons, India) and incubated at 28_°_C for 72 h without shaking.

### Biofilm inoculation with OS culture supernatant

20 µl of STM overnight culture and indicated volume (1% v/v or as indicated) of the OS culture supernatants were added to 2 ml of biofilm media and incubated under biofilm-promoting conditions(19). For co-culture experiments, the overnight cultures of the STM and OS or STM and PFA-fixed OS were added in a 1:1 ratio to 2 ml of biofilm media and incubated under biofilm-inducing conditions.

### Crystal Violet Staining

Post 72 h of biofilm induction, the media was discarded, and the plates were thoroughly washed in water. The plates were dried, and 2 ml of 1% Crystal Violet (CV) was added to each well and incubated for 15 min. Post 15 min incubation, the CV stain was removed, and the plate was thoroughly rinsed in water to remove excess stain. The stained biofilm was destained with 70% ethanol, and the intensity of the colour of the destained solution was quantified at OD_595_ in the Tecan plate reader (Infinite Pro 200) (19).

### Glass bead assay

1 mm glass beads were added onto the pellicle to assess the strength of the pellicle at the air-liquid interface. The initial weight of the glass beads was noted. The glass beads were added until the pellicle collapsed, and the final weight was noted down and plotted.

### Gut Permeability Assay

The infected mice cohorts were orally administered with FITC Dextran (25mg/kg of mice body weight, FD-4-100MG, Sigma) 4h before euthanization. The blood plasma was collected post-euthanization by cardiac puncture followed by centrifugation. The blood plasma was evaluated for mean fluorescence via plate reader (TECAN) and flow cytometry(CytoFlex, Beckman).

### MTT Assay

Post-16h treatment of pre-seeded Caco-2 cells with 10µg/ml of bacteriocin, MTT (3-(4,5-dimethylthiazol-2-yl)-2,5-diphenyltetrazolium bromide) staining solution (0.5mg/ml) was added to each well followed by incubation at 37°C CO_2_incubator for 4h. 50µl of DMSO was added per well for solubilizing the formazan crystals. Absorbance was measured at 570nm using a plate reader (TECAN).

### Transmission electron microscopy

Overnight STM or LM-grown cultures (in the presence or absence of OS supernatant or bacteriocin) were subcultured into fresh media (1:10) ratio and incubated at 37°C for 6 hours under shaking conditions. These samples were centrifugated at 1000rpm for 10 min to obtain the bacterial pellet. The bacterial pellet was subjected to PBS wash twice via centrifugation at 1000 rpm for 10 min.5 µl of the finally resuspended cell suspension was added on a glow-discharged copper-grid, air dried, stained with 1% uranyl acetate for 30 s, and visualized under a transmission electron microscope (JEM-100 CX II; JEOL).

### Supernatant Treatment

The supernatant was treated with 7.5 mM concentrations of EDTA and incubated at 65 ℃ for 1 h. 10 µl of proteinase-K (NEB, stock 20 mg/ml) was added to the 200 µl supernatant and incubated at 37 ℃ for 1 h. The proteinase was inactivated with 0.5 mM PMSF and incubated at 28 ℃ for 1 h. The supernatant was subjected to heating at 37_°_C (60 min), 65°C (15 min) or 95°C (15 min) and immediately frozen at − 20°C (19). Post-treatment, the supernatants were used to inoculate the biofilm to test the nature of the biofilm inhibitory molecules.

### Supernatant concentration and molecular weight-based separation

4 ml of the supernatants were transferred to the 3k MWCO or 10k MWCO Amicon ultra filter centrifugal device (Amicon® Ultra-4 centrifugal filter device, 3k MWCO, UFC800324; Amicon® Ultra-4 Centrifugal filter Unit, 10k MWCO, UFC801024) and centrifuged at 4000 g for 30 min in a swing bucket rotor(19). The concentrated solute from the bottom of the filter device and the flowthrough were collected and subjected to biofilm inoculation.

### Size Exclusion Chromatography

The bacterial supernatant was concentrated to 0.5ml using 3k MWCO-Amicon-Ultra-4-centrifugal unit and loaded onto the Superdex 200pg size exclusion column (XK16/60, 124mL; Cytiva, USA), pre-equilibrated Tris-Cl buffer (pH-8) containing 100mM NaCl and 20mM Tris-Cl. The sample was eluted with the same buffer at a flow rate of 0.8mL/min, and 1mL fractions were collected. The fractions corresponding to the elution peaks were analyzed for biofilm inhibition activity and subjected to mass spectrometry.

### LC/MS analysis of the supernatant fractions

The concentrated gel filtration fractions of supernatants were subjected to in-solution trypsin digestion for LC Q-TOF MS analysis. Briefly, 100 µg of total protein was resuspended in 100 µl of 7-8M of urea buffer. To this, 5 µl 200mM DTT was added and incubated at room temperature for 1 hr, followed by alkylation with 20µl of 200 mM iodoacetamide and incubation at room temperature for 1h in the dark. This was followed by the addition of 1mM of Calcium chloride and incubation at RT for 10 min. For protein digestion, trypsin solution was added to the sample to a final trypsin to protein ratio of 1:30 (w/w) and incubated at 37°C for 16 h, with frequent shaking. The reaction was stopped by adding 0.1% formic acid to adjust the pH to 3-4. The digested protein samples were desalted using a reverse phase C-18 column (Acclaim prep EASY-Spray (C18-RP)) using mobile phase solvent A-2% acetonitrile and solvent B- 80% acetonitrile. The sample injection volume was 10µl. All the samples were analyzed using Thermo Scientific SII for Xcalibur method. The MS scan was acquired using Orbitrap detector with 60000 resolution in the positive polarity and scan range (*m/z*) 350-2000. The ion source type was NSI with static voltage; positive ion (V)- 2200V and negative ion (V)-600 along with temperature settings at 275°C. This was followed by ddMS^2^ IT HCD with 30% fragmentation HCD collision energy using ion trap detector. The raw data was analyzed using Proteome Discoverer 2.5 against the *O. splanchnicus* proteome available in UniProt (Proteome ID: UP000006657).

### Purification of bacteriocin

*O. splanchnicus* culture supernatant was concentrated within the range of >3kDa and <10kDa using the Amicon Ultra-4 Centrifugal filter units of 10MWCO and 3MWCO. The concentrated cell-free supernatant containing the bacteriocin (∼7kDa) was precipitated by ammonium sulphate (80% to 98%). The precipitate was collected by centrifugation at 12000rpm for 30 min and dissolved in 1X PBS. The dissolved precipitate was again filtrated using the 3KDa cutoff Amicon centrifugal filter units. This concentrated fraction was further subjected to Fast Performance Liquid Chromatography using Superdex 75 Increase 10/300 GL- 24ml volume (GE Healthcare) column. The column was equilibrated with 1X Phosphate Buffered Saline buffer and 0.5ml of concentrated sample (23.6 mg/ml) was injected into the column. Based on the increase in Absorbance units at 280nm from baseline, 0.3ml fractions were collected at a retention volume of 13.3ml to 21.8ml (Fig. S8). The fractions were analyzed by 20% SDS-PAGE electrophoresis and stained with Coomassie staining solution.

### *In vitro* swim motility assay

2 µl (2*10^6^ CFU/ml) of treated or untreated stationary phase bacterial samples were spotted on the 0.3% agar (LB-STM, BHI-LM) plates supplemented with 0.5% yeast extract, 1% casein enzyme hydrolysate, 0.5% NaCl, and 0.5% glucose. The plates were incubated at 37°C and images were taken every 6 h (STM) or 24h (LM) using the camera of Biorad illuminator.

### Statistical analysis

Data were analyzed and graphed using the GraphPad Prism 8 software (San Diego, CA). Statistical significance was determined by Student’s t-test or One-way ANOVA followed by Dunnett’s Multiple Comparisons test to obtain p values. ANOVA was used during comparison among three or more groups whereas Student’s t-test was used to compare the difference between two groups. Mann Whitney t-test was used as a non-parametric counterpart of Student t-test when variances differed. The log-rank test also known as Mantel-Cox test is used to compare the survival percentages of the mice groups. Adjusted p-values below 0.05 are considered statistically significant (****p<0.0001, *** p < 0.001, ** p<0.01, * p<0.05). The results are expressed as mean ± SD or SEM of three independent experiments.

## RESULTS

### *O. splanchnicus* hinders *Salmonella* burden and its systemic dissemination

To denote the role of OS in *Salmonella* pathogenesis, we colonized C57BL/6 mice with 10^5^ CFU of OS 1 day prior to *Salmonella* infection via the oral route and validated the colonization by undertaking OS 16S rRNA qPCR analysis (Fig 1, A-B). OS pre-colonized mice showed decreased susceptibility to *Salmonella* infection in comparison to the non-colonized mice cohort which succumbed to death by day 9 post-infection (p.i.). Contrastingly, 100% of the OS colonized mice were alive even at day 11 post *Salmonella* infection (Fig 1, C). *Salmonella* colonization of the feces and intestinal tissue and dissemination to the MLN, blood, and liver were reduced upon OS pre-colonization (Fig 1, D-K). OS non-colonized mice exhibited increased weight loss percentage upon *Salmonella* infection and OS pre-colonization prevented weight loss (Fig 1, H). Our result suggests OS could ameliorate *Salmonella* colonization, organ burden and dissemination (Fig 1).

**Figure 1.**
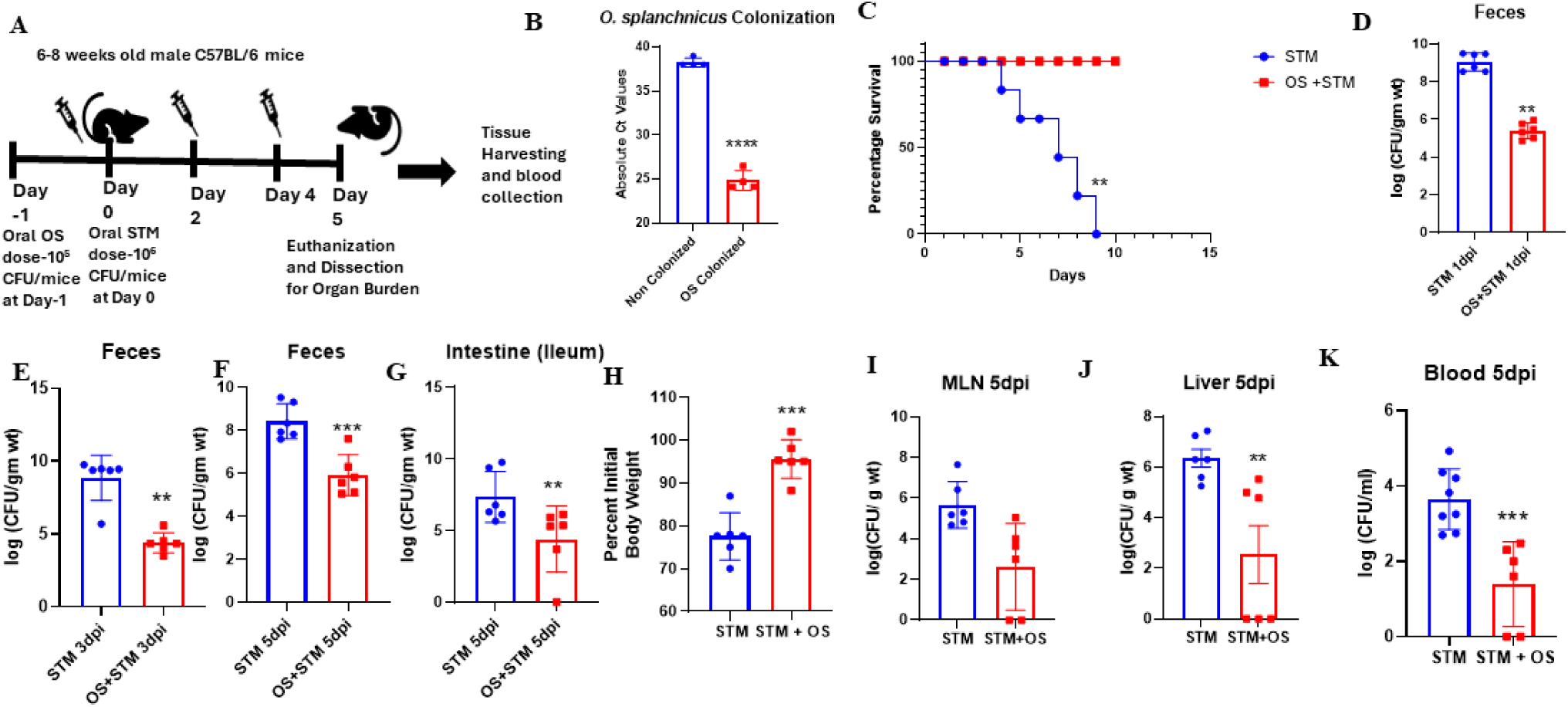
*Odoribacter splanchnicus* colonization protects mice from *Salmonella* colonization and dissemination. **A**-Schematic representation showing animal experimental strategy of infection. **B**-OS colonization validation via qPCR quantification of *O. splanchnicus* 16S rRNA from fecal samples of mice at 6^th^ day post-OS colonization . **C**-Survival percentage of non-colonized-STM-infected mice versus OS-colonized-STM-infected mice cohorts. Data is representative of N=3, n=5 where N denotes number of biological replicates and n denotes the number of mice per group. Mantel Cox or the log-rank test was used to determine the p values. **D-G**- STM organ loads in feces at 1^st^ day post-infection (D), 3^rd^ day post-infection (E), 5^th^ post-infection (F) and in ileal tissues (contents and wall) at 5^th^ day post-infection (G) in OS colonized and non-colonized mice cohorts. Data is representative of N=2, n=6 where N denotes number of biological replicates and n denotes the number of mice per group. Mann Whitney Test was performed to obtain p-values. **H**-Percent of initial weight (at Day 0) exhibited by OS colonized and non-colonized mice upon STM infection at 5^th^ day post-infection. Data is representative of N=3,n=6 where N denotes number of biological replicates and n denotes the number of mice per group. **I-K**- STM organ loads in MLN (I), liver (J) and blood (K) at 5^th^ post-infection in OS colonized and non-colonized mice cohorts. Data is representative of N=3, n=6 where N denotes number of biological replicates and n denotes the number of mice per group. Mann Whitney Test was performed to obtain p-values.

### *O. splanchnicus* colonization ameliorate the acute *Salmonella-induced* inflammation and its associated pathogenesis

Heightened inflammatory responses facilitate *Salmonella* dissemination in mice(20). Thus, we hypothesized that OS gut colonization might ameliorate the *Salmonella-*induced inflammation. ELISA-mediated IL-6 serum quantification revealed reduced IL-6 production upon *S.* Typhimurium (STM) infection in the OS colonized mice group to the non-colonized cohort (Fig 2, A). To combat the enteric pathogen challenge, the host employs one of its major defence arsenals, that is, the production of antimicrobial peptides (AMPs)(21, 22). STM infection induces enhanced mRNA expression of AMPs in mice with respect to the uninfected control as per previous literature findings (17). We found that OS colonization in mice prevented STM infection-induced expression of several AMP genes such as Lysozyme-C1 precursor, Angiogenin-4 precursor, CCL20, and Regenerating-islet-derived 3 beta-precursor (Fig 2, B). The haematoxylin eosin-stained intestinal sections suggested increased intestinal tissue damage with irregularly organized villi, necrotic submucosa layer, neutrophil infiltration, and detached intestinal cells in all the STM-infected non-OS-colonized cohorts. However, OS pre-colonization restored the intestinal tissue architecture in all the mice cohorts (Fig 2, C-D).

**Figure 2.**
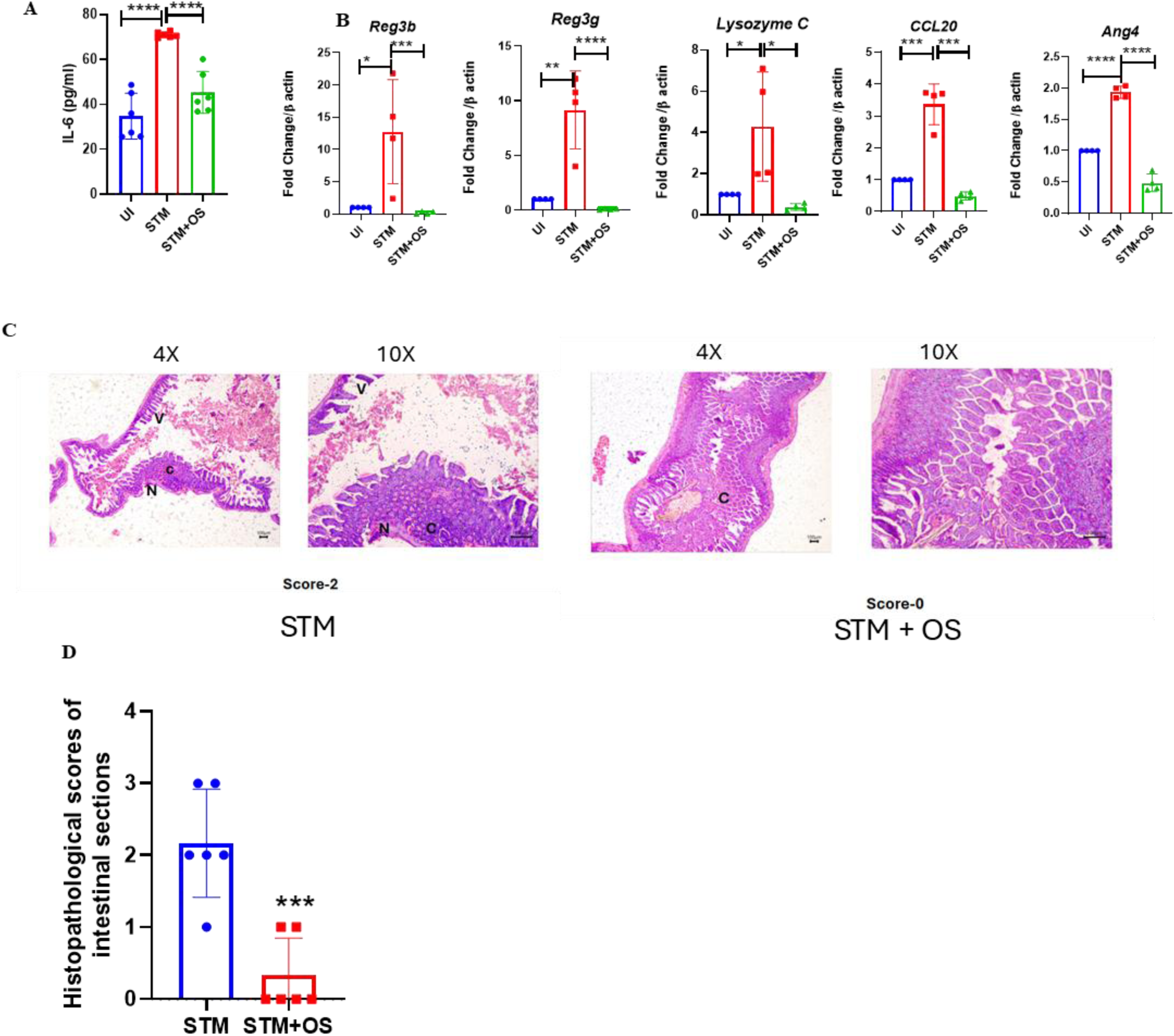
*O. splanchnicus* reduces *Salmonella-*induced inflammation and intestinal damage. **A**-Estimation of serum IL-6 production via ELISA upon STM infection in OS non-colonized and colonized mice cohorts at 5^th^ day post-*Salmonella* infection. Unpaired t-test is performed to obtain the p-values. Data is representative of N=3, n=3. **B**-Quantitative gene expression data of several antimicrobial peptides (AMPs) precursor genes in uninfected (UI) and STM-infected mice intestinal tissues harvested at 5^th^ dpi from OS colonized or non-colonized cohorts. Unpaired t-test is performed to obtain the p-values. Data is representative of N=3, n=4. **C**- **D**-Haematoxylin-eosin-stained intestinal sections harvested from STM-infected OS non-colonized or colonized mice cohorts at 5^th^ dpi. 0-represents normal pathology, 1-represents-mild pathology, 2-3-represents severe pathology.

### *O. splanchnicus* restricts *Salmonella-*mediated GVB damage and promotes tight junction gene expression

*Salmonella* breaches the gut vascular barrier (GVB) by inducing tight junction disruption. *Salmonella* like other food-borne antigens enters into the Mesenteric Lymph Node (MLN) by accessing the lymphatics and subsequently gains entry into the liver via the bloodstream(23). The translocation of the pathogen across the blood endothelial cells is governed by the integrity of the GVB(1). The GVB refers to the endothelial barrier and the microvasculature characterized by the elaborate framework of the junctional complexes (1). The GVB architecture is maintained by several tight junctions and adherens junction proteins that control paracellular trafficking(24, 25). Other cell types associated with the microvasculature of the GVB forming a vascular unit includes pericytes and fibroblasts(26). PV-1 (plasmalemma-vesicle associated protein-1) expression is a marker of GVB damage and heightened PV-1 expression denotes EC permeability(27, 28). Our data show reduced *PV-1* marker expression, alongside enhanced tight junction protein mRNA expression, such as *Tjp1* and *Claudin5*, upon OS colonization during *Salmonella* infection in comparison to the non-OS-colonized cohorts. (Fig. 3, A). Previous studies in *Mycobacterium marinum* have shown that transcriptional induction of pro-angiogenic markers like *Vegfa* contributes to increased bacterial burden and dissemination(29). In our study, OS pre-colonization in the gut resulted in reduced expression of the pro-angiogenic *Vegfa* in the infected liver tissues, which coincided with the reduced *S*. Typhimurium burden in the bloodstream and liver (Fig S1, A). Previous literature suggested reduced expression of *Axin2,* a β catenin target gene, upon wildtype *Salmonella* infection, even after Wnt3a stimulation suggesting that *Salmonella* affects the GVB by inhibiting canonical Wnt signalling (1). Here, our results suggest that OS pre-colonization restores the *Salmonella*-triggered attenuated Wnt signalling response with increased Axin2 expression in the *Salmonella*-infected intestinal tissues (Fig. S1, B). The integrity of the infected mice gut was estimated by the gut permeability assay using FITC-Dextran (Fig. 3, B). Our results indicate increased gut vascular permeability upon *Salmonella* challenge which declined upon OS pre-colonisation, implicating the ability of OS to prevent gut barrier disruption.

**Fig 3.**
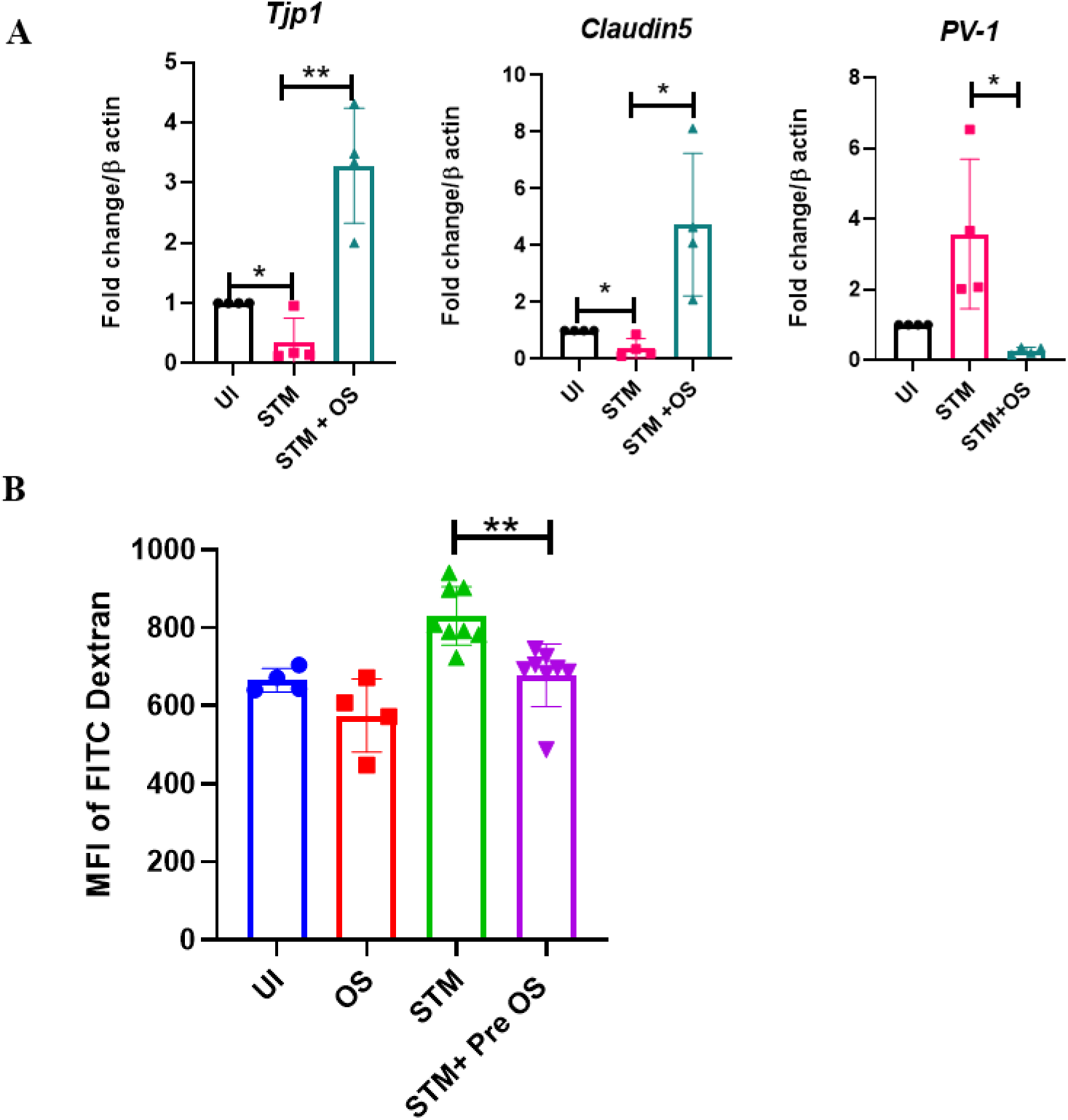
*Odoribacter splanchnicus* gut colonization prevents *Salmonella-*induced GVB damage and promotes tight junction gene expression. **A**-Quantitative gene expression studies of tight junction protein complexes (*Tjp1*, *Claudin5*) and PV-1 gene (marker of GVB damage) in STM-infected mice intestinal (ileal) tissue in presence or absence of OS gut colonization at 5^th^ day post-infection. Un-paired t-test was used to obtain the p-values. Data is representative of N=3, n=4. **B**- Estimation of FITC fluorescence in mice plasma via gut permeability assay in infected mice cohorts upon oral administration of FITC Dextran (25mg/kg) 4h before euthanization at 5^th^ day post *Salmonella* infection. Un-paired t-test was used to obtain the p-values. Data is representative of N=4.

### *O. splanchnicus* culture supernatant restricts *Salmonella* biofilm formation and intracellular proliferation in human intestinal cells

To delineate the mechanism behind the protective role of OS against *Salmonella* pathogenesis, we performed *Salmonella* infection studies in human intestinal epithelial cell line Caco-2 in the presence of OS. The intracellular survival assay suggested decreased STM invasion and subsequently attenuated intracellular replication within the infected Caco-2 cells upon co-infection with live OS and in the presence of OS culture supernatant (Fig 4, A-B). However, STM co-infection with fixed OS showed percent invasion and intracellular replication like that of STM infection (Fig 4, A-B). Together, these results hinted at the presence of some secreted inhibitory components in the OS culture supernatant that has the propensity to limit STM infection. The reduced invasion and proliferation of STM within Caco-2 cells in presence of OS was subsequently validated in immunofluorescence studies, wherein STM showed a significant reduction in invasion at 2hr p.i. and intracellular proliferation at 16hr p.i. in case of OS co-infection (Fig 4, C-D). Similar results were obtained upon infection with the human-restricted serovar STY to the Caco-2 cells wherein OS and its supernatant attenuated STY invasion and intracellular replication (Fig. 4, E-F). Further, we assessed the *in vitro* growth of the STM in co-culture conditions with OS or in the presence of OS culture supernatant in anaerobic conditions to mimic the microaerophilic or hypoxic environment of the gut or in aerobic culture conditions. We observed no significant delay *in vitro* growth of STM at its initial lag to log phase of its growth in the presence of OS supernatant or with live OS under anaerobic culture conditions (Fig S2, A-B). In aerobic growth conditions as well, *Salmonella* growth remained unperturbed even in the presence of OS or its culture supernatant (Fig S2, C-D). Several previous studies have corroborated the ability of commensal bacterial culture supernatants to inhibit colonization and the biofilm formation of several pathogens. For instance, a commensal bacterium, *Staphylococcus epidermis*, could inhibit biofilm formation and nasal colonization of its pathogenic counterpart, *Staphylococcus aureus* (30). Moreover, biofilm formation is also regulated at the interspecies level. Indole is an inter-species biofilm regulator that decreases *E. coli* biofilm formation and facilitates biofilm formation of pseudomonads (31). STM forms biofilm to colonize the host, evade host defence arsenals and antibiotics, persist during chronic infections, and transmit to new hosts (32, 33). We, therefore, examined the biofilm pellicle strength and biofilm formation of STM via crystal violet staining on the solid-liquid interface in the presence or absence of live, PFA-fixed-OS or OS culture supernatant by inoculating STM in biofilm-inducing media with low salinity. STM exhibited biofilm formation defect in the presence of live OS or OS supernatant (Fig 4, G). However, the STM biofilm inhibitory property of OS was diminished upon treatment with paraformaldehyde (PFA)-fixed OS (Fig 4, G). Further, qPCR-mediated gene expression studies of *Salmonella* biofilm genes revealed decreased expression of genes regulating biofilm (*csgD*, *csgA*, *bapA*) under treatment with live OS or its culture supernatant (Fig.4, H-J). Flagella-mediated attachment and several virulence factors such as SPI-1 and SPI-2 genes are responsible for *Salmonella* pathogenesis and its biofilm formation (19, 34, 35). Therefore, we evaluated a few of the *Salmonella* flagellar and SPI-1 and SPI-2 gene expression in the presence or absence of OS or its culture supernatant via quantitative real-time PCR. Our results depicted attenuated expression of flagellar gene (*fliC*), SPI-1 (*sipC*) and SPI-2 (*pipB2*) genes both in the presence of live OS and its spent media but not in presence of PFA fixed-OS (Fig. 4, K-M). The loss in *Salmonella* flagellar motility was further validated via TEM imaging wherein OS or its supernatant reduced the flagella formation of *Salmonella* considerably (Fig.4, N). Together, our results implicate the ability of OS to inhibit STM biofilm formation via active secretion of certain inhibitory molecules in the supernatant.

**Fig 4.**
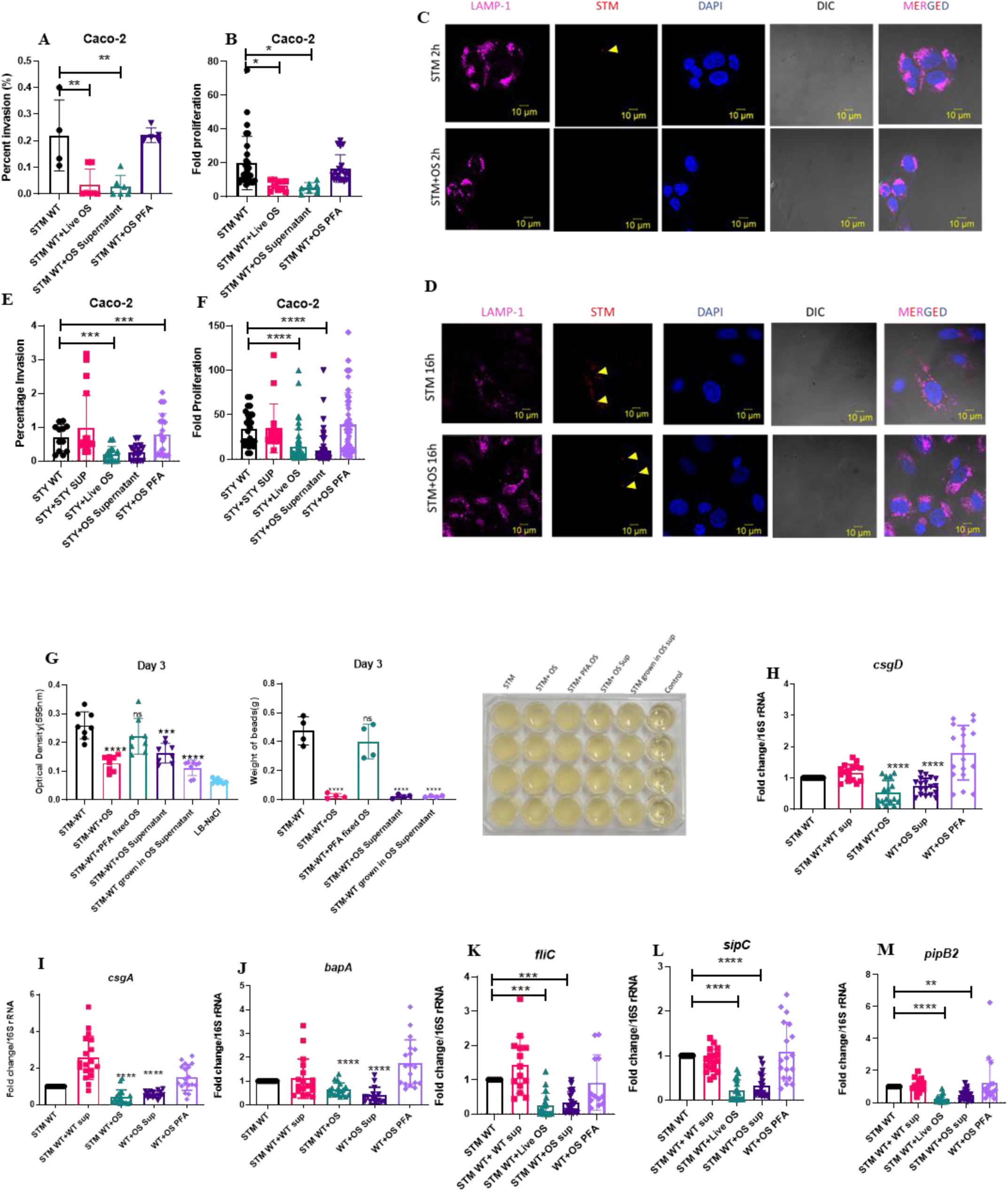
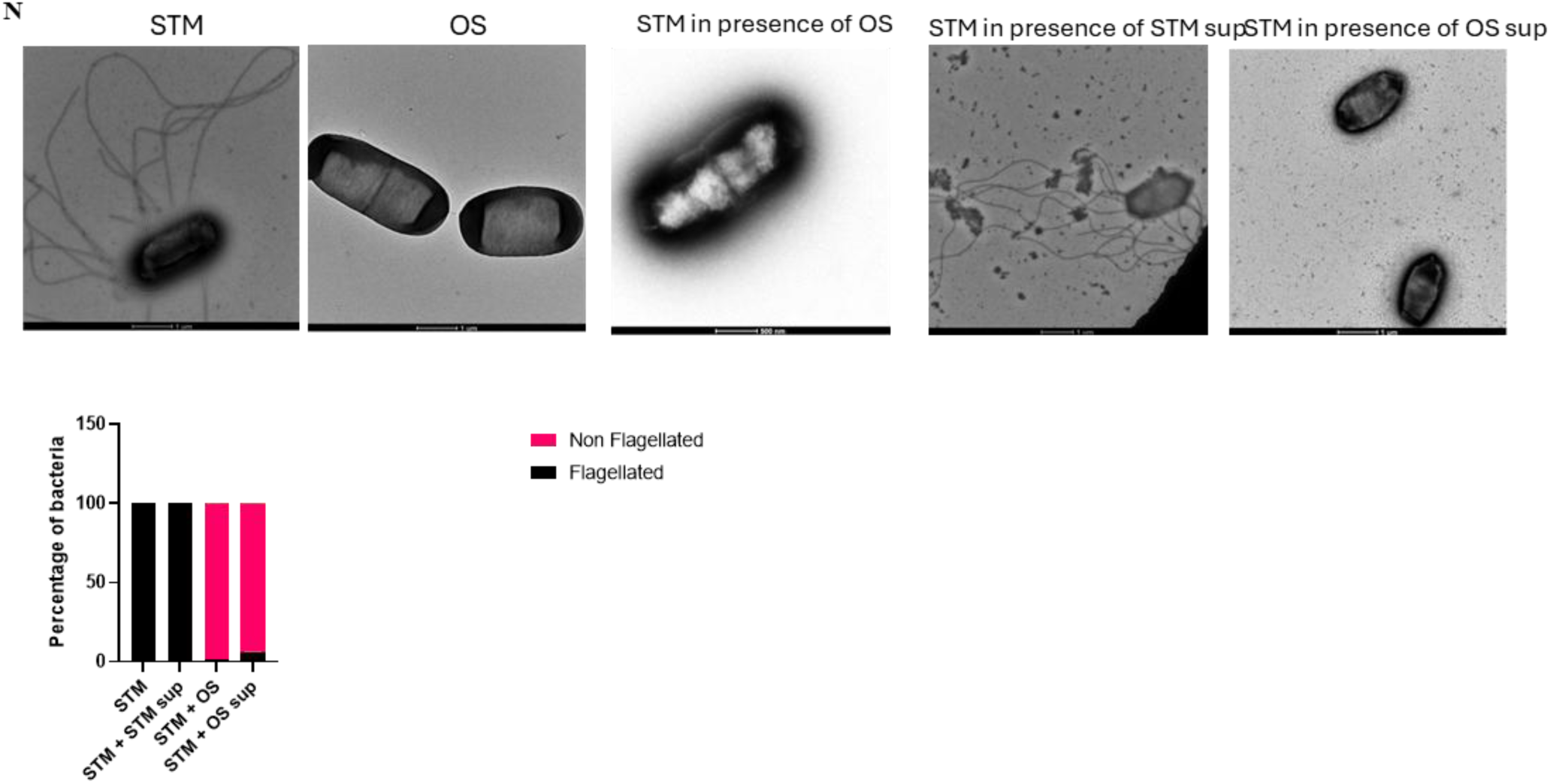
*O. splanchnicus* exhibits its inhibitory effect on *Salmonella* biofilm formation and inhibits invasion and intracellular replication in human intestinal cells. **A**-Percent invasion of STM in human Caco-2 human intestinal cells in presence or absence of live OS, OS culture supernatant, PFA-fixed-OS. Data is representative of N=3, n=4. One-way ANOVA was used to determine the p-values. **B**-Intracellular proliferation of STM Caco-2 cells in presence or absence of live OS, OS culture supernatant, PFA-fixed-OS. Data is representative of N=3, n=4. One-way ANOVA was used to determine the p-values. **C**- Representative immunofluorescence images depicting STM invasion into Caco-2 cells in the presence or absence of OS at 2hr post-infection. Data is representative of N=3, n=100. **D**- Representative immunofluorescence images depicting STM invasion into Caco-2 cells in presence or absence of OS at 2hr post-infection. Data is representative of N=3, n=100. **E**-Percent invasion of STY in Caco-2 human cells in presence or absence of OS, OS culture supernatant, PFA-fixed OS. Data is representative of N=3, n≥9. One-way ANOVA was used to determine the p-values. **F**-Intracellular proliferation of STY in Caco-2 cells in presence or absence of live OS, OS culture supernatant, PFA-fixed-OS. Data is representative of N=3, n≥9. One-way ANOVA was used to determine the p-values. **G**- Biofilm assay showing the biofilm formation ability of STM-WT or STM-WT grown in OS supernatant after 72h of inoculation in presence or absence of live OS, OS culture supernatant, PFA-fixed-OS. Data is representative of N=4, n=4.One-way ANOVA was used to determine the p-values. Image of the STM biofilm-formed plate in the above stated conditions has been shown. **H-M-** RT-qPCR mediated gene expression of STM biofilm regulatory genes (*csgD*) (H), *csgA* (I), *bapA* (J), along with flagellar gene (*fliC*) (K), SPI-1 (*sipC*) (L) and SPI-2 (*pipB2*) (M) genes in presence or absence of live OS or its spent media or PFA-fixed-OS. Data is representative of N=4, n=4. One-way ANOVA was used to determine the p-values. **N-** Transmission Electron Microscopic (TEM) images of STM in presence or absence of *Salmonella* WT supernatant, OS and its supernatant to assess the possession of flagella. The quantification plot exhibits the percentage of bacterial flagellated or non-flagellated cells (number of cells quantified n≥ 100)

### *O. splanchnicus* inhibitory effect on *Salmonella* is specific as *Enterococcus faecalis*, another gut resident microbe, fail to inhibit *Salmonella* intracellular proliferation or biofilm formation

The resident gut microbes provide colonization resistance against invading pathogens by deploying several mechanisms, including the production of inhibitory molecules, nutrient competition, maintenance of the intestinal epithelial integrity, immune activation, and via the deployment of bacteriophages (36, 37). To understand whether our observed inhibitory trait of OS against *Salmonella* is due to a common colonization restriction phenomenon of gut microbes via competitive inhibition, or it is a specific phenomenon, we evaluated the role of another gut resident microbe, *Enterococcus faecalis* on *Salmonella* intracellular replication in human intestinal epithelial cells and *Salmonella* biofilm formation. *Enterococcus sp.* belong to the phylum of Firmicutes in the family Enterococcaceae. Approximately, 10^6^-10^7^ *Enterococcus sp*. reside in the human intestine with major abundance of *E. faecalis* (10^5^-10^7^ CFU/gm feces) and *E*. *faecium* (10^4^-10^5^/gm feces) (38). *E. faecalis*, *E. faecium*, *Bifidobacterium sp*. *Lactobacillus sp.,* and *E. coli* colonize the gut of healthy breastfed infants during the initial 7-10 days post birth (39). Our intracellular survival assay data showed no defect in *Salmonella* invasion or intracellular replication capability in Caco-2 cells in presence or absence of *E. faecalis*, its culture supernatant or PFA-fixed *E. faecalis* (Fig. S3, A-B). Moreover, *E. faecalis* or its *E. faecalis* supernatant did not inhibit *Salmonella* biofilm formation (Fig. S3, C). Together, our results suggest the inability of *E. faecalis* to inhibit *Salmonella* invasion, intracellular proliferation or its *in vitro* biofilm formation, thereby highlighting the specificity of OS in restricting *Salmonella* pathogenesis.

### *O. splanchnicus* confers protection against another foodborne pathogen *Listeria monocytogenes*

To understand whether OS exerts a common broad-spectrum protection or colonization resistance against other foodborne pathogens such as *Listeria monocytogenes*, we orally infected 6-8-week-old C57Bl/6 mice with 10^8^ CFU of *L. monocytogenes* (LM) 1 day post colonization with 10^5^ CFU of *O. splanchnicus* for organ burden evaluation via oral gavage. For survival study, mice were orally gavaged with 10^9^ CFU of *L. monocytogenes*. OS*-*colonized mice exhibited increased survival rates upon LM challenge compared to the non-colonized mice cohort (Fig. 5, A). In line with the survival data, the increased bacterial burden was observed in the blood, feces, intestinal tissue, MLN, spleen and liver alongside enhanced percent loss in mice body weight in the case of *Listeria* infection in non-colonized mice group with respect to OS-precolonized- mice group (Fig. 5, B-J). In a similar line to *Salmonella* infection, OS mitigated host antimicrobial response with reduced expression of antimicrobial peptide transcripts during LM infection, amounting to reduced intestinal tissue damage. Further, OS promoted tight junction functions during *Listeria* infection with reduced expression of GVB damage marker, *PV-1* and angiogenesis marker, *Vegf* expression (Fig. S4, A-C). Moreover, OS colonization could restrict the *in vivo Salmonella* and *Listeria* biofilm-like microcolonies within the spleens of the infected mice cohorts as opposed to the non-colonized mice cohorts (Fig.S4, D-E). We estimated the communities of bacterial cells within the spleen tissue via Calcofluor white staining (40). Calcofluor white stains the biofilm matrix or extracellular polymeric substances (EPS) comprised of cellulose (40, 41) and exopolysaccharides that help in the adherence of the bacterial cells. This is the first study demonstrating biofilm-like bacterial colonies of *Salmonella* or *Listeria* secreting EPS in spleens. Like *Salmonella*, *in vitro* culture of *Listeria* with OS or its culture supernatant led to a loss in flagella as evidenced via the TEM imaging (Fig.S4, F). Since flagella is one of the factors facilitating biofilm formation and invasion, OS could inhibit both *Salmonella* and *Listeria* colonisation and biofilm formation. Further, LM exhibited attenuated intracellular proliferation within human intestinal Caco-2 cells upon live OS or OS supernatant treatment (Fig. 6). Altogether, this result hints at the potency of OS to act broadly against Gram-positive and Gram-negative bacteria.

**Fig 5.**
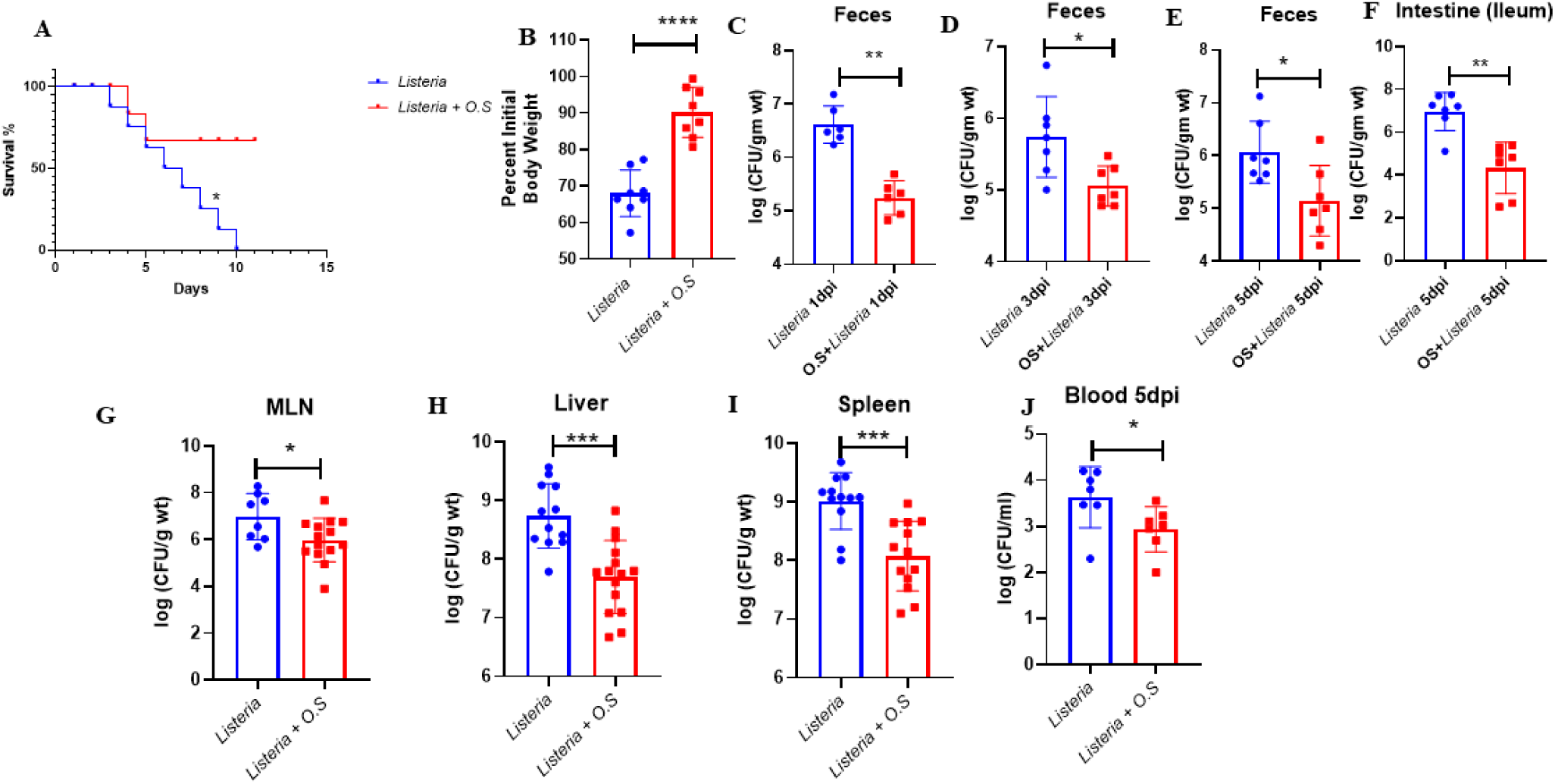
*Odoribacter splanchnicus* exerts its protective role against *Listeria monocytogenes* in mice. **A**- Survival percentage of non-colonized LM-infected mice versus OS-colonized *Listeria*-infected mice cohorts. Data is represantative of N=2, n=8, where N denotes number of biological replicates and n denotes the number of mice per group. Mantel Cox or the log-rank test was used to determine the p values. **B**-Percent of initial weight (at Day 0) exhibited by OS colonized and non-colonized mice upon LM infection at 5^th^ dpi. Data is representative of N=2,n=8, where N denotes number of biological replicates and n denotes the number of mice per group. Mann Whitney Test was performed to obtain p-values. **C-F**- LM organ loads in feces at 1^st^ day post-infection (C), 3^rd^ day post-infection (D), 5^th^ post-infection (E) and in ileal tissues (contents and wall) at 5^th^ day post-infection (F) in OS pre-colonized and non-colonized mice cohorts. Data is representative of N=2, n=7 where N denotes number of biological replicates and n denotes the number of mice per group. Mann Whitney Test was performed to obtain p-values. **G-J**- LM organ loads in Mesenteric lymph Node (MLN) (G), liver (H), spleen (I) and blood (J) at 5^th^ dp. in OS colonized and non-colonized mice cohorts Data is representative of N=2, n=8, where N denotes number of biological replicates and n denotes the number of mice per group. Mann Whitney Test was performed to obtain p-values.

**Fig. 6.**
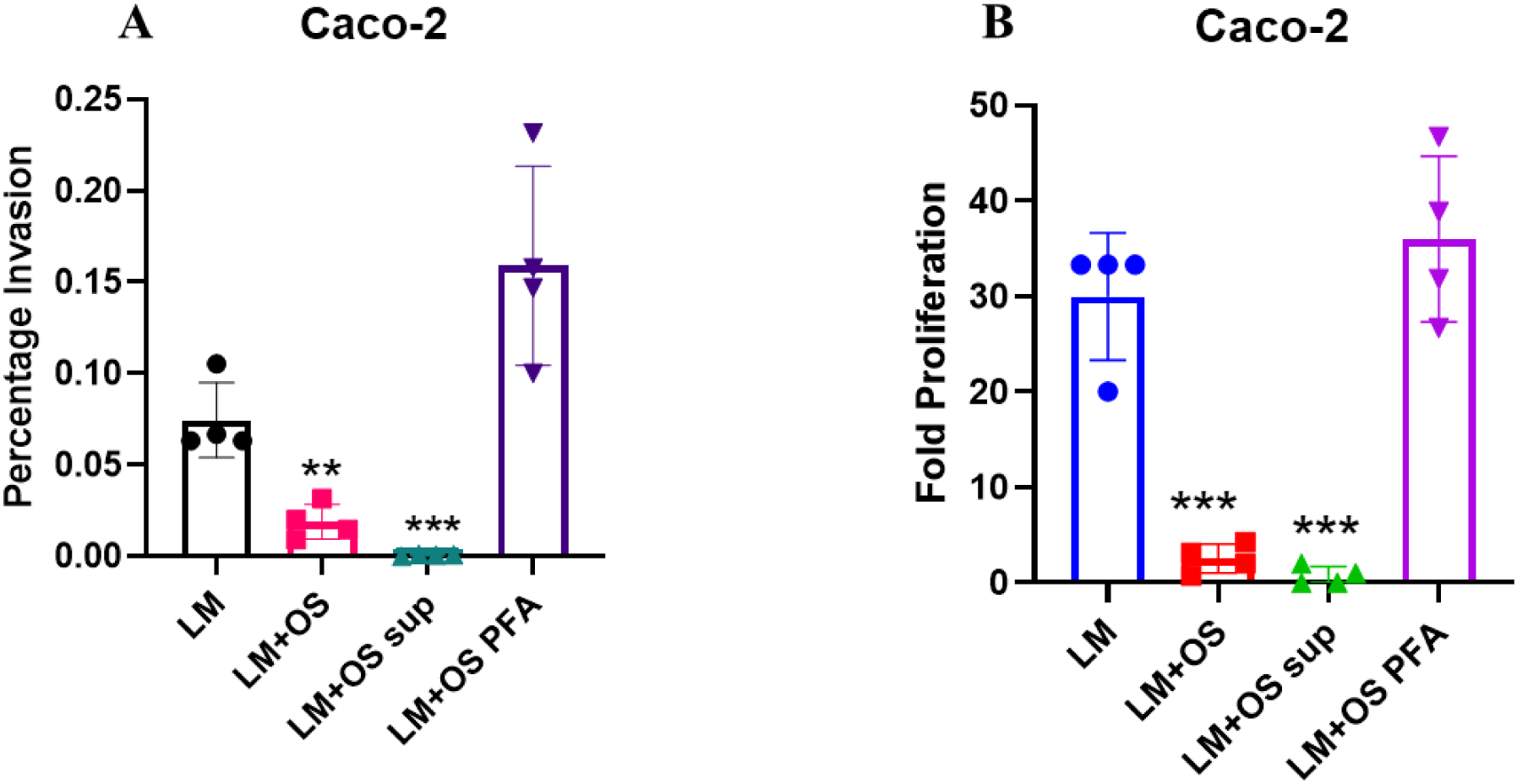
O*d*oribacter *splanchnicus* and its culture supernatant restricts *Listeria monocytogenes* pathogenesis in human intestinal cells. **A**- Percent invasion of LM in Caco-2 cells in presence or absence of live OS, OS sup, PFA-fixed-OS. Data is representative of N=3, n=4. One-way ANOVA and Multiple Comparison test are used to obtain the p-values. **B**-Intracellular fold proliferation of LM in Caco-2 cells in the presence or absence of live OS, OS sup, PFA-fixed-OS. Data is representative of N=3, n=4. One-way ANOVA and Multiple Comparison test are used to obtain the p-values.

### The protective role of *O. splanchnicus* against *Salmonella* Typhimurium and *Listeria monocytogenes* persists even on administration post-establishment of infection

Next, we sought to examine whether the protective role of OS against *Salmonella* and *Listeria* is sustained when the infection has been fully developed and whether it can serve as a treatment regimen against foodborne diseases. Our results indicate the protective ability of OS against both foodborne pathogens in their intestinal phase of infection when administered at 1-day post-infection (dpi) as depicted by lower pathogen loads in all the organs and blood along with reduced percent weight loss at 5th-day post-*Salmonella* or *Listeria* challenge in the OS-administered (1-dpi) cohort when compared to the untreated cohorts (Fig. 7,A-K, Fig. S5, A-C). Although the protective ability of OS persists even upon establishment of infection when administered at 3rd-day post-*Salmonella* or *Listeria* challenge as indicated by reduced organ loads and percent weight loss in the OS-colonized cohorts, the extent of protection is lower compared to its administration at 1-dpi (Fig. 7, L-W, Fig. S5, D-S), Further, the OS challenge at 1-dpi reduced the mice’s susceptibility to the *Salmonella* and *Listeria* challenge(Fig. 7, L,R). However, administration of OS at 3-dpi delayed the death of the *Salmonella* and *Listeria*-infected mice cohorts by 1 day (Fig. 7, L,R). In line with the survival and organ burden data, OS treatment at both 1^st^ and 3^rd^ dpi reduced both *Salmonella* and *Listeria*-induced GVB damage with reduced PV-1 expression, AMPs, and *Vegf* expression, alongside promoting tight junction protein (*Tjp1* and *Claudin5*) and *Axin2* expression (Fig. S5, D-M). However, OS administration at 3-dpi conferred less protection than at 1-dpi as indicated by the protein expression profiles of PV-1, and tight junction protein ZO-1 (Fig. S5, N-O). Additionally, OS colonization mitigated gut vascular permeability during *Salmonella* and *Listeria* challenges as indicated by reduced FITC-dextran fluorescence units in the plasma both upon pre or post-OS colonization (Fig. S5, P). Moreover, the hematoxylin- eosin-stained intestinal sections revealed reduced intestinal tissue damage with restoration in the villi architecture in the 1- and 3-dpi OS-administered mice cohorts compared to the non-OS-treated groups (Fig.S5, Q). Moreover, OS colonization could restrict the *in vivo Salmonella* and *Listeria* biofilm within the spleens of the infected mice cohorts as opposed to the non-colonized mice cohorts both when administered at 1-day and 3-day post-pathogen challenge. (Fig.S5, R-S).

**Fig. 7.**
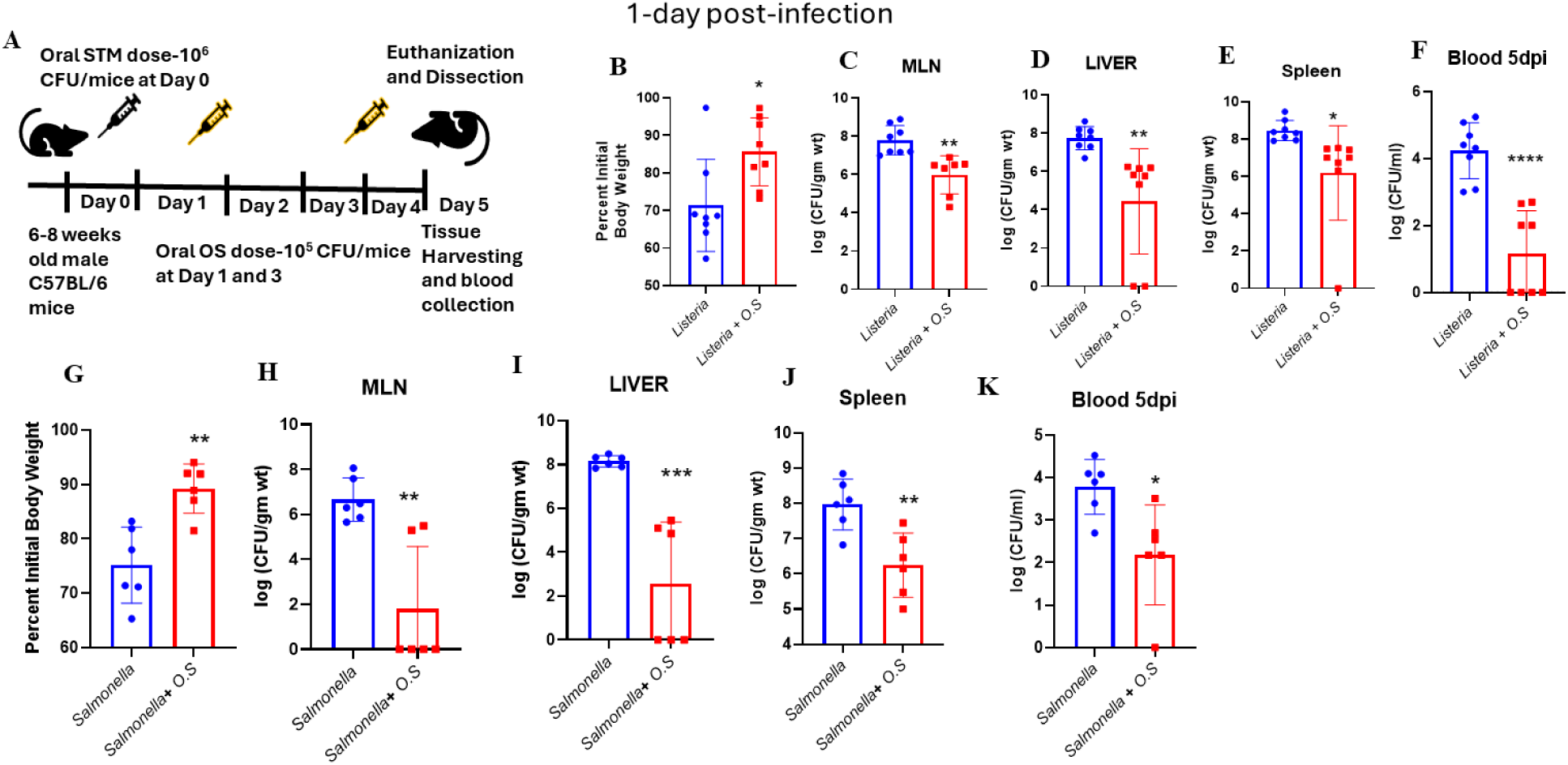
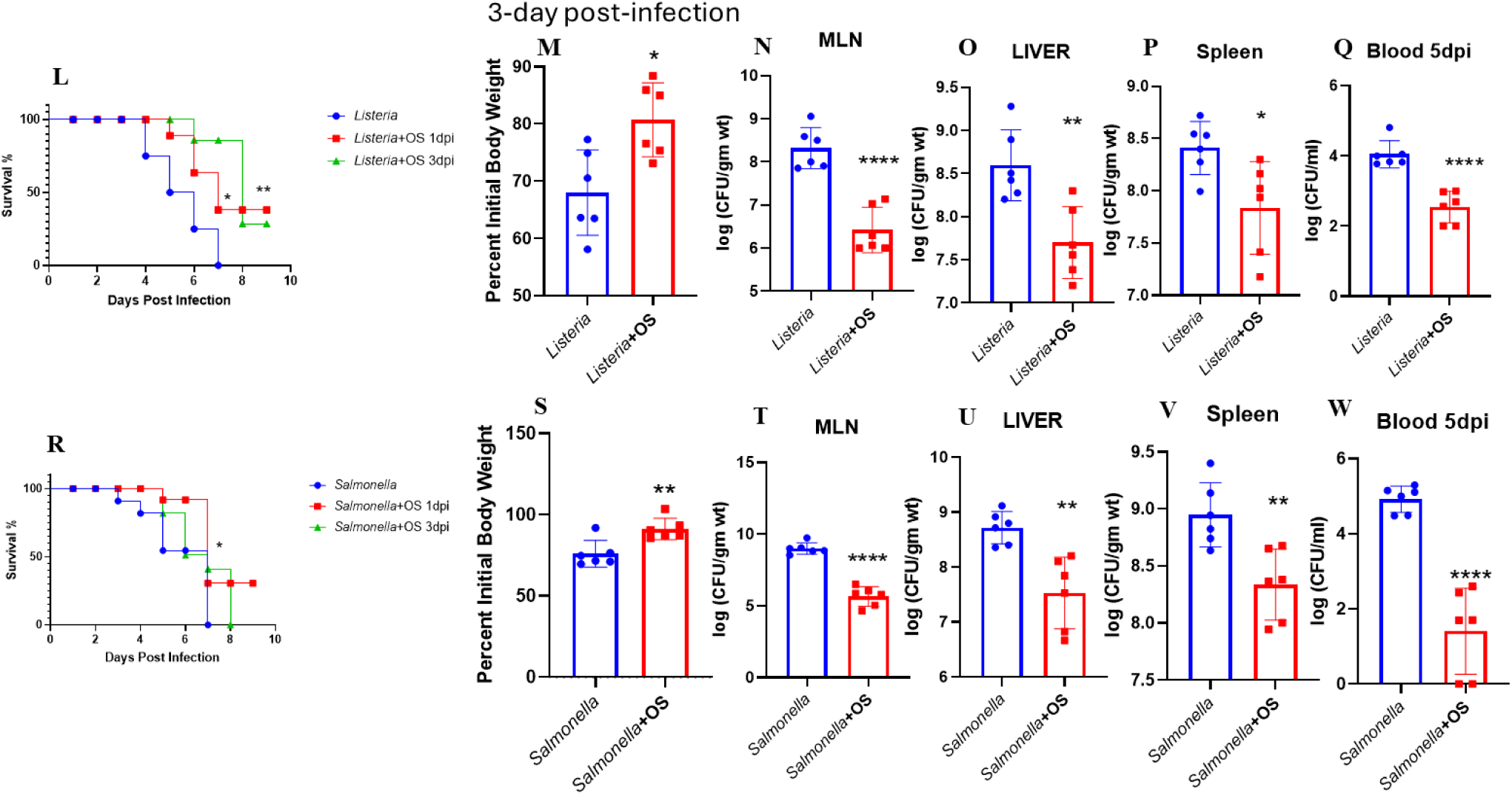
*O. splanchnicus* even confers protection against foodborne pathogens when administered post-pathogenic challenge. **A-** Schematic representation showing the animal experimental strategy of infection. **B-** Percent of initial weight (at Day 0) exhibited by OS colonized (1-dpi) and non-colonized mice upon LM infection at 5^th^ dpi. Data is representative of N=2,n=8, where N denotes number of biological replicates and n denotes the number of mice per group. Mann Whitney Test was performed to obtain p-values. **C-F**- LM organ loads in MLN (C), liver (D), spleen (E) and blood (F) at 5^th^ dpi upon OS administration at 1-dpi. Data is representative of N=2, n=8, where N denotes number of biological replicates and n denotes the number of mice per group. Mann Whitney Test was performed to obtain p-values. **G**- Percent of initial weight (at Day 0) exhibited by OS colonized (1-dpi) and non-colonized mice upon STM infection at 5^th^ dpi. Data is representative of N=2,n=8, where N denotes number of biological replicates and n denotes the number of mice per group. Mann Whitney Test was performed to obtain p-values. **H-K**- STM organ loads in MLN (H), liver (I), spleen (J) and blood (K) at 5^th^-dpi upon OS administration at 1-dpi. Data is representative of N=2, n=8, where N denotes number of biological replicates and n denotes the number of mice per group. Mann-Whitney Test was performed to obtain p-values. **L**- Survival percentage of non-colonized LM*-*infected mice versus OS-colonized (1 dpi or 3 dpi) *Listeria-*infected mice cohorts. Data is representative of N=2, n=8, where N denotes number of biological replicates and n denotes the number of mice per group. Mantel Cox or the log-rank test was used to determine the p values. **M**- Percent weight loss exhibited by OS colonized (3-dpi) and non-colonized mice upon LM infection at 5th-day post-infection. Data is representative of N=2,n=8, where N denotes number of biological replicates and n denotes the number of mice per group.Mann-Whitney Test was performed to obtain p-values. **N-Q**- LM organ loads in MLN (N), liver (O), spleen (P) and blood (Q) at 5^th^-dpi upon OS administration at 3-dpi. Data is representative of N=2, n=8, , where N denotes number of biological replicates and n denotes the number of mice per group. Mann-Whitney Test was performed to obtain p-values. **R**- Survival percentage of non-colonized *Salmonella-*infected mice versus OS-colonized (1 dpi or 3 dpi) *Listeria-*infected mice cohorts. Data is represantative of N=2, n=8, where N denotes number of biological replicates and n denotes the number of mice per group. Mantel Cox or the log-rank test was used to determine the p values. **S**- Percent weight loss exhibited by OS-colonized (3-dpi) and non-colonized mice upon STM infection at 5th-day post-infection. Data is representative of N=2,n=8, where N denotes number of biological replicates and n denotes the number of mice per group. Mann-Whitney Test was performed to obtain p-values. **T-W**-STM organ loads in MLN (T), liver (U), spleen (V), and blood (W) at 5^th^-dpi upon OS administration at 3-dpi. Data is representative of N=2, n=8, where N denotes number of biological replicates and n denotes the number of mice per group. Mann-Whitney Test was performed to obtain p-values.

### Characterization of the inhibitory active molecule within *O. splanchnicus* culture supernatant

Our results suggested that not only live OS but also its spent culture supernatant could inhibit *Salmonella* invasion and intracellular proliferation in intestinal human epithelial cells. Moreover, the OS and its supernatant could inhibit STM biofilm formation *in vitro.* This inhibitory effect disappeared upon co-culture with paraformaldehyde-fixed-OS indicating the presumable role of certain inhibitory microbial products secreted by metabolically live OS. To delineate the nature of the secreted molecule/s, we subjected the supernatant to various conditions such as high temperature, divalent cationic chelator, EDTA, and protease (proteinase K), and subsequently assessed the biofilm formation (Fig. 8, A-C). Our biofilm assay suggested that the active molecule is resistant to EDTA treatment and is stable at higher temperatures (Fig. 8, A-B). Nevertheless, the active molecules within the fraction were sensitive to the proteinase K treatment, suggesting the possibility of being protein in nature (Fig. 8, C). SDS-PAGE of the OS supernatant revealed the size of the differential proteins with respect to the media control ranged below 65kDa. (Fig. 8, D). Concentration-dependent estimation of the biofilm activity of the supernatant revealed that the active molecules retain their activity at a minimal concentration of 0.4% v/v (Fig. 8, E). We further fractionated the supernatant using an Amicon 3k MWCO and 10k MWCO ultra filter device (Fig. 8, F-G). We found that the active molecule/s is/ are higher than 3kDa cut-off range eliminating the role of small molecules and ions (Fig. 8, F-G). A summary of the conditions tested for active molecule characterisation and their effective outcome has been summarized in Fig. 8, H. Subsequently, we subjected the culture supernatant to gel filtration chromatography (Fig. S6, A). Out of all the fractions tested, around 24 fractions showed a biofilm inhibitory role. These 24 fractions were further subjected to a titration experiment to zero down upon the three fractions with the highest inhibitory activity (A82,A83,A84) (S6, B-D). The gel filtration standard curve run suggested that the active molecules fall in the size range of <60KDA and >3kDa (Fig. S6, A). These three fractions along with the total concentrated supernatant (serving as positive control) and fractions with no or least inhibitory activity (serving as negative control-A31,B12) were ultimately subjected to Reverse Phase LC-MS mass spectrometry. Upon analysis of the proteomics dataset (Fig. S6, E), we obtained 364 proteins showing enrichment in the positive fractions. Among the 364 proteins, 74 proteins exhibited known or predicted function as per UniProt and STRING database and 11 proteins were linked to antibacterial, bacteriostatic, flagella, and biofilm regulation based on data curation and literature survey (Table-S1). We selected bacteriocin proteins exhibiting direct association with antibacterial activity for further analysis. These proteins include bacteriocins (A0A412WSU3 and A0A412WSS0) which are antimicrobial peptides that can kill or inhibit closely related or unrelated bacterial strains(42, 43). We purified the bacteriocins via ammonium sulphate precipitation followed by gel filtration chromatography to obtain a single fraction comprising of bacteriocins with close-range molecular weight (6.6 kDa and 7.156kDa, respectively) (Fig. S7).

**Fig. 8.**
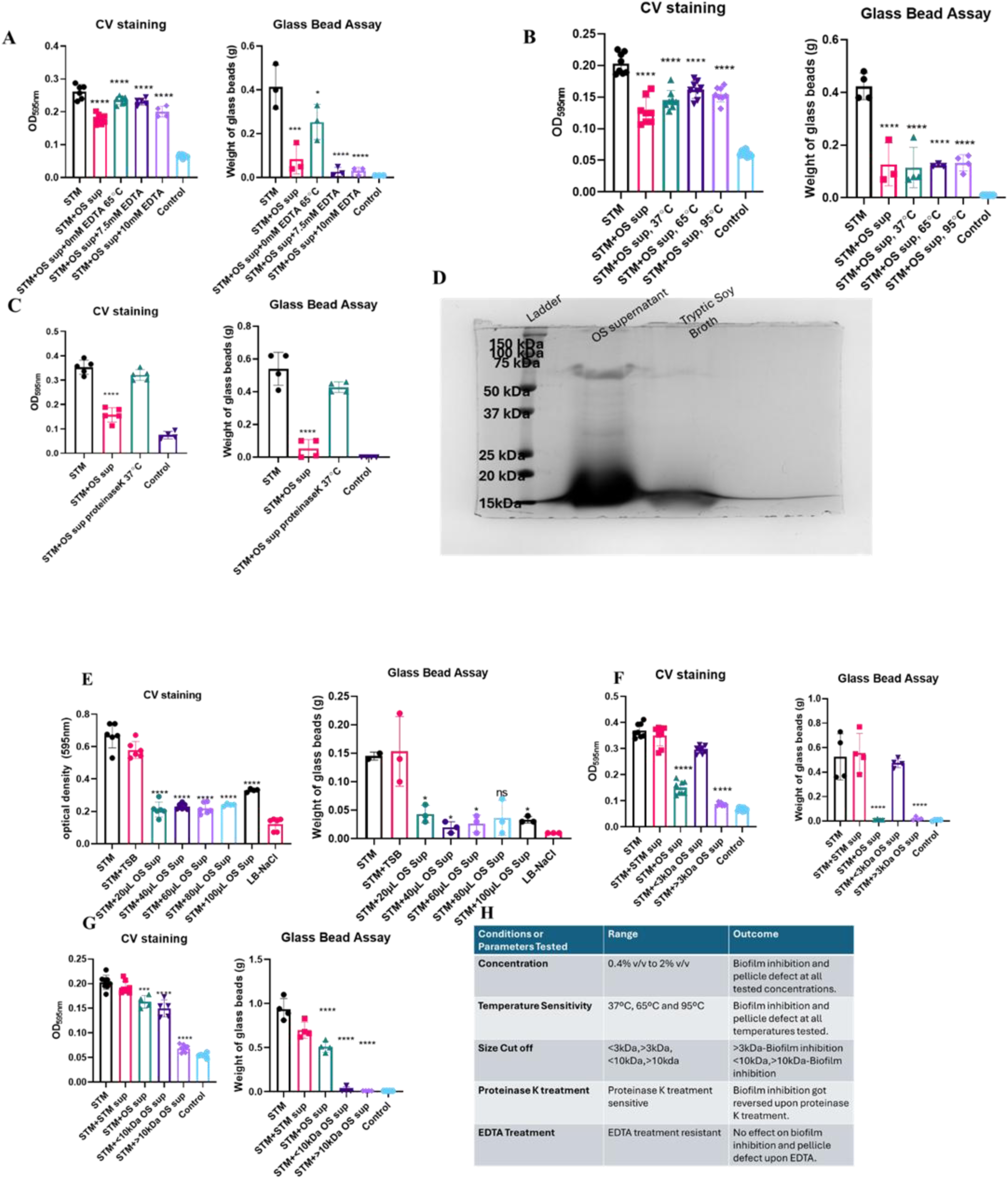
Characterization of the nature of inhibitory active molecule within *O. splanchnicus* culture supernatant. **A-** Biofilm assay showing the biofilm formation ability of STM after 72h of inoculation in presence of OS culture supernatant upon EDTA treatment. Data is representative of N=4, n=4. One-way ANOVA is used to obtain the p-values. Statistical test is performed with respect to STM. **B**- Biofilm assay showing the biofilm formation ability of STM after 72h of inoculation in presence of OS culture supernatant at high temperature conditions. Data is representative of N=4, n=4. One-way ANOVA is used to obtain the p-values. Statistical test is performed with respect to STM. **C-** Biofilm assay showing the biofilm formation ability of STM after 72h of inoculation in presence of OS culture supernatant under Proteinase K treatment. Data is representative of N=4, n=4. One-way ANOVA is used to obtain the p-values. Statistical test is performed with respect to STM. **D**- Coomassie-stained SDS-PAGE demonstrating the size range of the differential protein bands present in OS supernatant as opposed to the tryptic soy broth media control. **E**- Biofilm assay showing the biofilm formation ability of STM after 72h of inoculation in presence of varying concentration OS culture supernatant. Data is representative of N=4, n=4. One-way ANOVA is used to obtain the p-values. Statistical test is performed with respect to STM. **F**- Biofilm assay showing the biofilm formation ability of STM after 72h of inoculation in presence of OS culture supernatant fractions fractionated using 3kDa Amicon filter. Data is representative of N=4, n=4. One-way ANOVA is used to obtain the p-values. Statistical test is performed with respect to STM. **G**- Biofilm assay showing the biofilm formation ability of STM after 72h of inoculation in presence of OS culture supernatant fractions fractionated using 10kDa Amicon filter. Data is representative of N=4, n=4. One-way ANOVA is used to obtain the p-values. Statistical test is performed with respect to STM. **H**- Summary table demonstrating the conditions tested for active molecule characterization and their effective outcome.

### *O. splanchnicus*-derived purified peptide, bacteriocin, limits both *Salmonella* and *Listeria* pathogenesis *in vitro*

Treatment of the Caco-2 cells with the purified bacteriocin inhibited STM, STY, and LM invasion and fold proliferation without compromising host-cell viability (Fig. 9, A-D). Additionally, the purified bacteriocin treatment inhibited STM biofilm formation in a dose-dependent manner alongside reduced transcript level expression of *Salmonella* biofilm regulatory genes (Fig. S8, A-D). Moreover, bacteriocin treatment (10µg/ml) attenuated *Salmonella* virulence gene expression such as SPI-1 gene- *sipC*, SPI-2 gene-*pipB2,* and flagellar gene *fliC* (Fig. S8, E-G). Subsequently, TEM imaging revealed a complete loss of *Listeria* flagella upon treatment with the OS-derived bacteriocin in contrast to *Salmonella* which retained few flagella even on bacteriocin treatment (Fig. 9, E-H). Furthermore, the motility assay demonstrated a significant swimming defect of *Listeria* and *Salmonella* upon treatment with either OS supernatant or bacteriocin to the untreated control (Fig. 9, I-J). Overall, our study depicts the novel role of OS-derived bacteriocin in limiting *Salmonella* and *Listeria* invasion and pathogenesis by regulating virulence effectors such as SPI-1, SPI-2 genes, biofilm regulatory genes, and flagella.

**Fig. 9.**
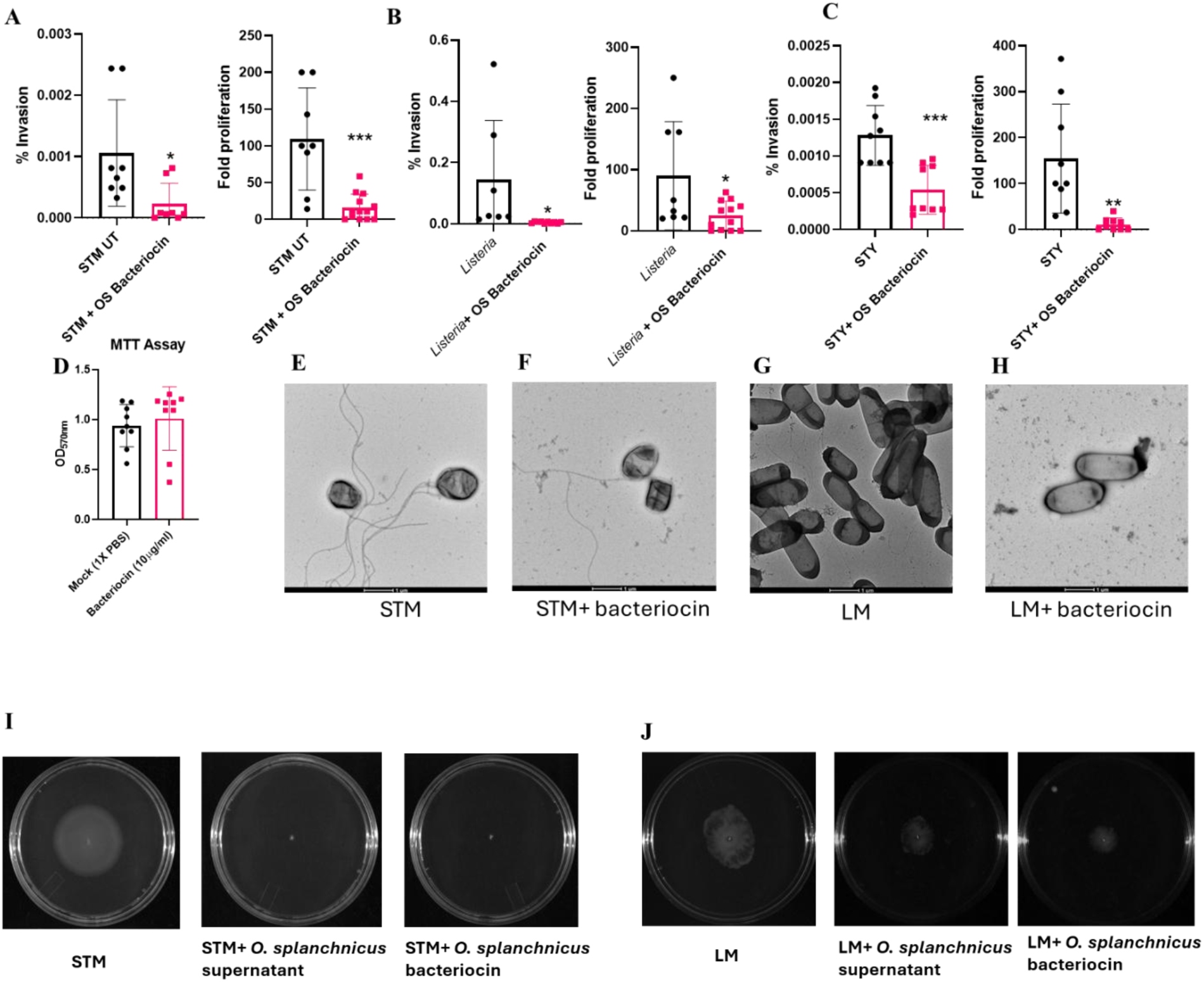
*O. splanchnicus* bacteriocin restricts both *Salmonella* and *Listeria* pathogenesis *in vitro*. **A**- Percent invasion and intracellular fold proliferation of STM in Caco-2 cells under OS bacteriocin (10µg/ml) treatment. Data is representative of N=2, n=3. Unpaired t-test is used to obtain the p-values. **B**- Percent invasion and intracellular fold proliferation of LM in Caco-2 cells under OS bacteriocin (10µg/ml) treatment. Data is representative of N=2, n=3. Unpaired t-test is used to obtain the p-values. **C**- Percent invasion and intracellular fold proliferation of STY in Caco-2 cells under OS bacteriocin (10µg/ml) treatment. Data is representative of N=2, n=3. Unpaired t-test is used to obtain the p-values. **D**-MTT assay evaluating the viability of Caco-2 cells upon 16h of mock or bacteriocin (10µg/ml) treatment. Data is representative of N=2, n=4. **E-F**-TEM images of STM under untreated (D) or bacteriocin (10µg/ml) (E) treatment to assess the possession of flagella. **G-H**- TEM images of LM under untreated (D) or bacteriocin (10µg/ml) (E) treatment to assess the possession of flagella. **I-** Swimming motility assay of STM under OS supernatant or bacteriocin treatment. **J**- Swimming motility assay of LM under OS supernatant or bacteriocin treatment.

## DISCUSSION

*Salmonella* gains access to the systemic sites by traversing the gut intestinal barrier via utilizing several routes. Its primary site of entry is the distal ileum wherein it induces the SPI-1 Type Three Secretion System (T3SS) effector proteins to facilitate its uptake. After its initial entry into the enterocytes and M-cells overlying the Peyer’s patches, *Salmonella* colonizes the draining MLN and eventually disseminates to the liver and spleen via the circulating lymphatics and the bloodstream (44). *Salmonella* uptake by M cells facilitates its stealthy entry via avoidance of the mucus barrier. Additionally, the presence of flagella allows them to propel through the viscous mucus and activate the immune responses(45). The onset of systemic dissemination coincides with GVB disruption and loss of tight-junction protein expression (1). Several reports have suggested the protective role of gut commensals in maintaining gut epithelial health against food-borne pathogen insults. *E. coli* provide colonization resistance against STM in the presence of a specific microbial consortium by limiting galactitol wherein *Lachnospiraceae* consume the free sugar, conferring colonization resistance (46). Moreover, *E. coli* contribute to colonization resistance by competing with *S.* Enteritidis for aerobic respiration (47). Similarly, the commensal *Klebsiella michiganensis* retention or transmission provided colonization resistance to Enterobacteriaceae by exhibiting nutrient competition (48).

Previous literature hinted toward the beneficial effects of OS in healthy human gut wherein reduced OS abundance have been linked to obese gut signature (49). Furthermore, OS colonization in mice have been shown to enhance Foxp3^+^/RORγt^+^ regulatory T cells, interleukin IL-10 and short chain fatty acids production, limiting colitis in mouse models (50). OS have been depicted to confer protection against colitis and colorectal cancer in mice by inducing intestinal Th17 development(13).

In this study, we report the broad-spectrum protective role of the gut commensal of bacteroidales order *O. splanchnicus* that limits *Salmonella* and *Listeria* invasion, pathogenesis, and systemic dissemination by promoting colonization resistance and regulating virulence effectors such as SPI-1, SPI-2, flagella, and biofilm regulatory genes. Additionally, not only OS but its culture supernatant could exert its inhibitory effect on *Salmonella* invasion and intracellular proliferation in intestinal human epithelial cells. Moreover, the OS and its supernatant could inhibit STM biofilm formation by regulating flagella formation. This STM biofilm inhibitory effect was however lost upon co-culture with paraformaldehyde-fixed-OS indicating the presumable role of certain inhibitory microbial products secreted by metabolically live OS. Further, our studies identified OS-produced bacteriocins responsible for restricting intracellular replication of *Salmonella* and *Listeria* in intestinal epithelial cells by regulating bacterial virulence arsenals such as biofilm regulatory genes, SPI-1, SPI-2 genes, and flagellar machinery. However, the presence of flagella on *Salmonella* post-bacteriocin treatment suggests the potential involvement of additional secreted accessory molecules within the OS supernatant and indicates that bacteriocins might act in concert with additional factors through diverse mechanisms, warranting future research. Bacteriocin-mediated suppression of several virulence-associated phenotypes with minimal impact on overall bacterial growth suggests that the bacteriocin may interfere with regulatory pathways controlling virulence. One possible mechanism is the modulation of global regulatory networks that coordinate virulence gene expression, potentially through subtle physiological perturbations or cellular signalling pathways. Such physiological changes could attenuate virulence programs including motility required for host cell invasion and intracellular survival. Alternatively, bacteriocin exposure may induce stress responses that indirectly downregulate energy-intensive virulence processes. However, determining the precise molecular mechanisms underlying these effects will require dedicated mechanistic studies, such as transcriptomic, proteomic, or regulatory pathway analyses, which are beyond the scope of the present study but warrant further investigation in future work.

Although the bacteriocin demonstrated significant anti-virulence activity in vitro, its proteinaceous nature presents potential challenges for oral therapeutic applications due to susceptibility to degradation in the gastrointestinal environment. Consequently, oral administration experiments in mice were not pursued in this study. Future work should therefore explore strategies to enhance stability and delivery, such as encapsulation and protective formulations. Addressing these challenges will be important for translating bacteriocin-based approaches into practical therapeutic applications.

Previous studies have suggested that bacteriocin expression by commensal bacteria can influence gut niche competition and bacteriocin delivery can serve as an effective therapeutic strategy to eliminate colonization of multi-drug resistant bacteria in the gastrointestinal tract without disrupting the indigenous microbiota (51, 52). Several studies have highlighted the role of commensal-produced bacteriocins. Bacteriocin-producing *Streptococcus salivarius* has been shown to effectively target the CRC-causing pathogen *Fusobacterium nucleatum* in a model of the human distal colon(53). *Lactobacillus plantarum* bacteriocin has been linked to intestinal and systemic improvements in diet-induced obese mice and confers epithelial barrier integrity (54). Further studies show that a class IIa bacteriocin, Pediocin PA-1, has an anti-bacterial effect against *L. monocytogenes* (55).

Bacteriocins produced by commensal bacteria are typically characterized by strong bactericidal activity that directly inhibits the growth of competing microorganisms. In contrast, the OS-derived bacteriocin described in this study primarily suppressed virulence-associated phenotypes including motility, SPI-1 and SPI-2 gene expression, and intracellular proliferation without markedly impairing bacterial growth. This observation suggests that, rather than acting solely as a classical antimicrobial compound, the OS-bacteriocin may interfere with regulatory pathways controlling virulence. One possible mechanism is the modulation of global regulatory networks that coordinate virulence gene expression, potentially through subtle physiological perturbations or cellular signalling pathways. Alternatively, bacteriocin exposure may induce stress responses that indirectly downregulate energy-intensive virulence processes. However, determining the precise molecular mechanisms underlying these effects will require dedicated mechanistic studies, such as transcriptomic, proteomic, or regulatory pathway analyses, which are beyond the scope of the present study but warrant further investigation in the future. OS-derived bacteriocin’s virulence-modulating activity is conceptually distinct from many previously described commensal-derived bacteriocins that mainly exert growth-inhibitory effects. The ability to attenuate virulence while exerting limited pressure on bacterial viability may represent a potentially advantageous strategy, as anti-virulence approaches are thought to impose reduced selective pressure for resistance compared with conventional bactericidal agents. Nevertheless, further mechanistic studies will be required to determine whether this effect arises from direct interference with virulence regulatory networks or from broader physiological perturbations induced by bacteriocin exposure.

This study provides evidence about inter-species regulation of gut colonization in maintaining gut homeostasis and protecting against several invading foodborne pathogens. Moreover, the long-term protective ability conferred by OS during pre-or post-infection colonization to mice against broad-spectrum foodborne pathogens such as *Salmonella* and *Listeria* highlights its therapeutic potential as a prebiotic for prophylaxis or post-infection treatment option, paving the way for future therapeutic interventions to thwart food-borne infections.

## ACKNOWLEDGEMENTS

We thank Nikhita Kirthivasan for her support during the commencement of the project. We also thank our Divisional Mass Spectrometry facility, TEM facility, Departmental Confocal Facility, Divisional Flowcytometry Facility and Central Animal Facility at IISc.

## AUTHOR CONTRIBUTION

Conceptualization: DH, DC

Methodology: DH, DM, RV, VD (Gel Filtration Chromatography), DD (Fast Performance Liquid Chromatography)

Investigation: DH, DM, RV, VD (Gel Filtration Chromatography), DD (Size exclusion chromatography for bacteriocin purification), RSR (Scoring of haemoxylin and eosin-stained samples)

Visualization: DH, DM, RV Formal analysis: DH Funding acquisition: DC

Project administration: DH, DC

Supervision: DC, UT (Proteomics), TH (Size Exclusion Chromatography), MG (Size Exclusion Chromatography)

Writing – original draft: DH

Writing – review & editing: DH, DM, RV, RSR, VD, DD, TH, MG, UT, DC

## DATA AND MATERIALS AVAILABILITY

All data are available in the main text or the supplementary materials.

The article has been submitted to the preprint server (56).

## SUPPLEMENTARY FIGURES AND LEGENDS

**Fig S1.**
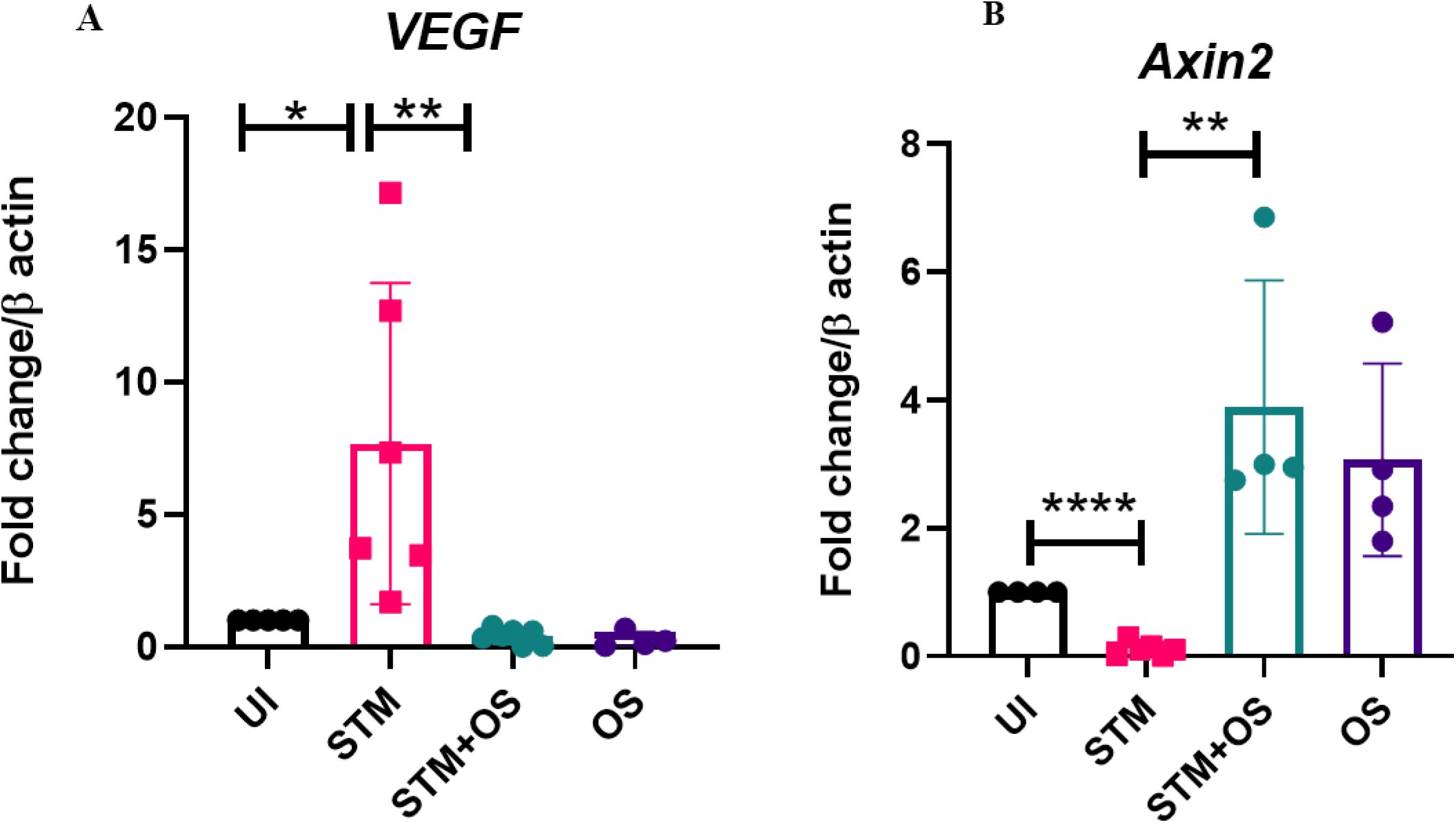
*O. splanchnicus* gut colonization prevents *Salmonella* dissemination by restricting intestinal GVB damage and angiogenesis. **A**- qPCR mediated gene expression of pro-angiogenic marker *Vegf* in STM-infected mice liver tissues in presence or absence of OS pre-colonization. Data is representative of N=2, n=4. Un-paired t-test was used to obtain the p-values. **B**- qPCR mediated gene expression of *Axin2* in STM-infected mice intestinal tissues in the presence or absence of OS pre-colonization. Data is representative of N=2, n=4. Un-paired t-test was used to obtain the p-values.

**Fig. S2.**
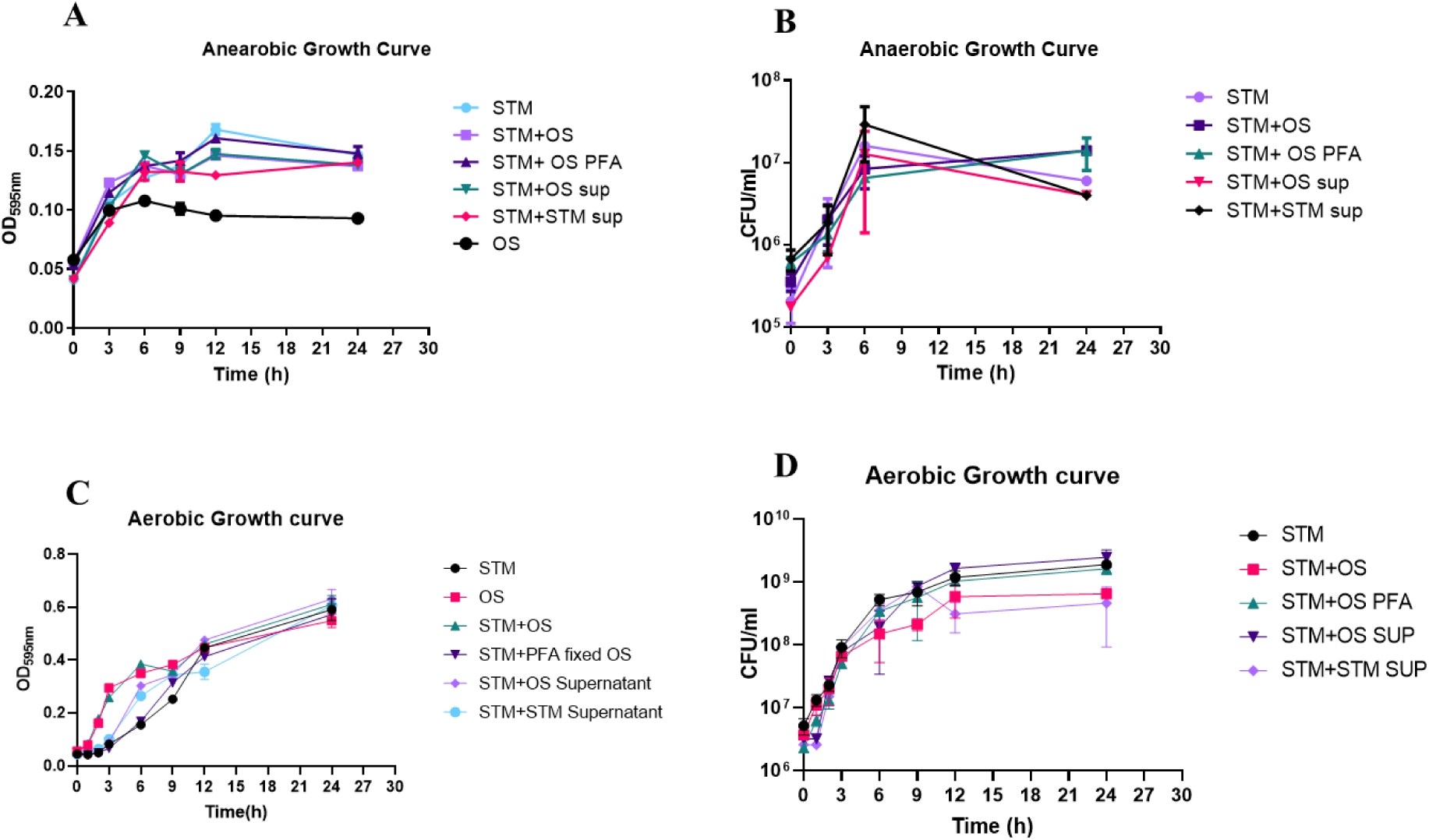
In vitro growth kinetics of *Salmonella* Typhimurium in presence of live or dead *O. splanchnicus* or its culture supernatant. **A-B** Growth kinetics of STM-WT, OS, STM-WT in presence of OS, STM-WT in presence of STM or OS culture supernatant and STM-WT in presence of PFA-fixed OS in LB under anaerobic growth conditions. **C-D**- Growth kinetics of STM-WT, OS, STM-WT in presence of OS, STM-WT in presence of STM or OS culture supernatant, STM-WT supernatant, and STM-WT in presence of PFA-fixed OS in LB under aerobic growth conditions.

**Fig S3.**
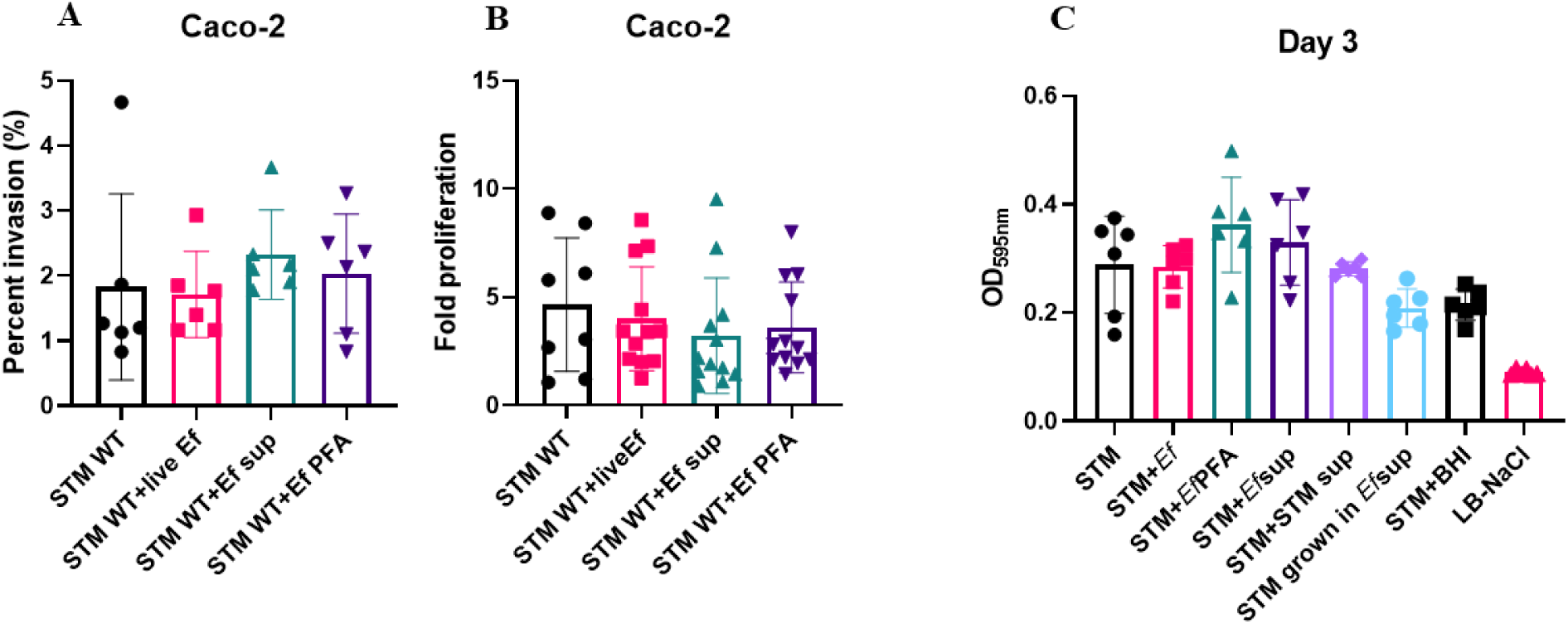
Unlike *O. splanchnicus, E. faecalis* fail to exert any inhibitory effect on *Salmonella in vitro* biofilm formation or *Salmonella* invasion and intracellular replication in human intestinal cells. **A**-Percent invasion of STM in Caco-2 cells in presence or absence of live *E. faecalis* (Ef), Ef culture supernatant, PFA-fixed-Ef. Data is representative of N=3, n=3. One-way ANOVA is used to obtain the p-values. **B**-Intracellular proliferation of STM in Caco-2 cells in presence or absence of live Ef, Ef culture supernatant, PFA-fixed-Ef. Data is representative of N=3, n=3. One-way ANOVA is used to obtain the p-values. **C**- Biofilm assay depicting the biofilm formation ability of STM-WT or STM-WT grown in Ef supernatant after 72h of inoculation in presence or absence of live Ef, Ef or STM culture supernatant, PFA-fixed. Crystal-violet staining was performed to assess the biofilm formation on the solid-liquid interphase and the [OD]_595_ absorbance reading was recorded. Data is representative of N=3, n=3. One-way Anova is used to obtain the p-values.

**Fig. S4.**
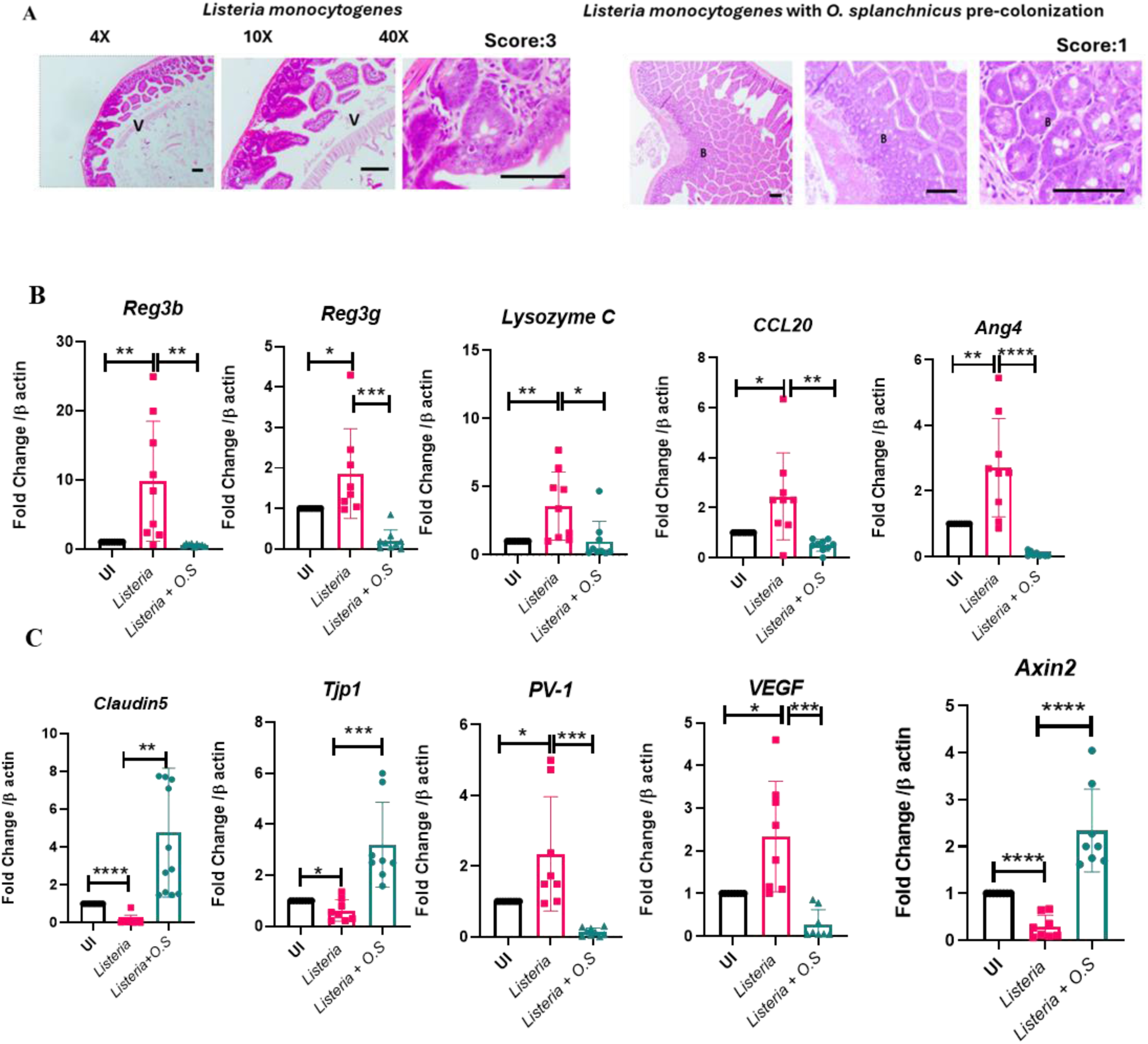
*Odoribacter splanchnicus* gut colonization protects mice against *Listeria monocytogenes* pathogenesis similar to *Salmonella* infection. **A**-Haematoxylin-eosin-stained intestinal sections harvested from OS non-colonized or colonized mice cohorts at 5^th^-day-post*-*LM-infection. 0-represents normal pathology, 1-represents- mild pathology, 2-3-represents severe pathology. **B**-Quantitative gene expression data of several antimicrobial peptides (AMPs) precursor genes in LM-infected mice intestinal tissues harvested at 5^th^ day post-infection from OS colonized or non-colonized cohorts. Unpaired Student’s t-test is performed to obtain the p-values. Data is representative of N=2, n=9. **C**- Quantitative gene expression study of tight junction protein genes *Tjp1*, *Claudin5*, GVB damage marker *PV-1*, angiogenesis marker-*Vegf,* and *Axin2* in LM- infected mice intestinal tissues harvested at 5^th^-dpi from OS colonized or non-colonized cohorts. Unpaired Student’s t-test is performed to obtain the p-values. Data is representative of N=2, n=5. **D-**Representative Calcofluor white (CW) staining of STM-infected mice spleen tissues at 5^th^-dpi. Data is representative of N=2, n=8 (tissue samples). Un-paired t-test was used to obtain the p-values. **E**- Representative CW staining of LM-infected mice spleen tissues at 5^th^-dpi. Data is representative of N=2, n=8 (tissue samples). Un-paired t-test was used to obtain the p-values. **F**- Transmission Electron Microscopic (TEM) images of LM in presence or absence of LM supernatant, OS and its supernatant to assess the possession of flagella. The quantification plot exhibits the percentage of bacterial flagellated or non-flagellated cells (number of cells quantified n≥ 100)

**Fig. S5.**
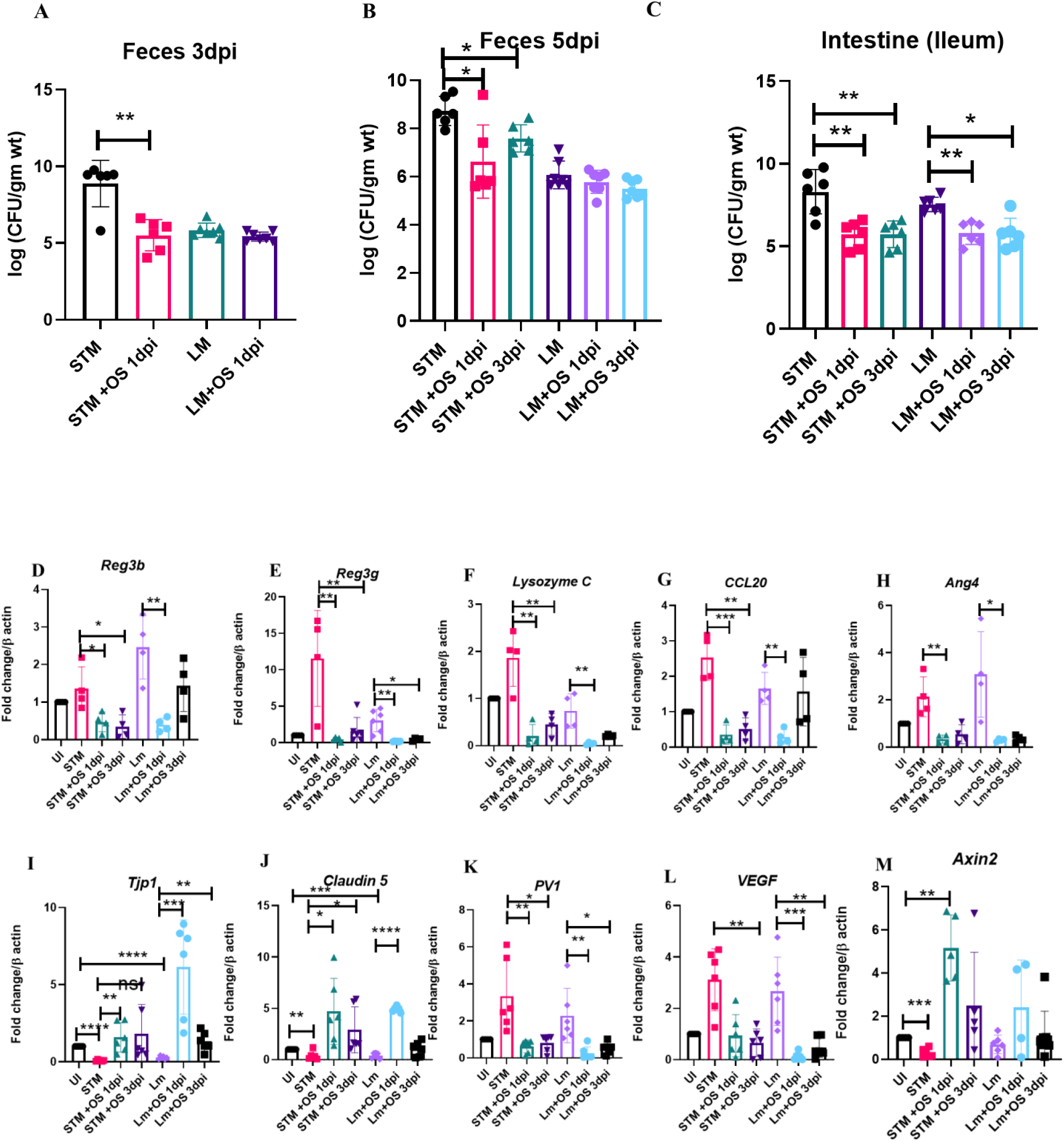
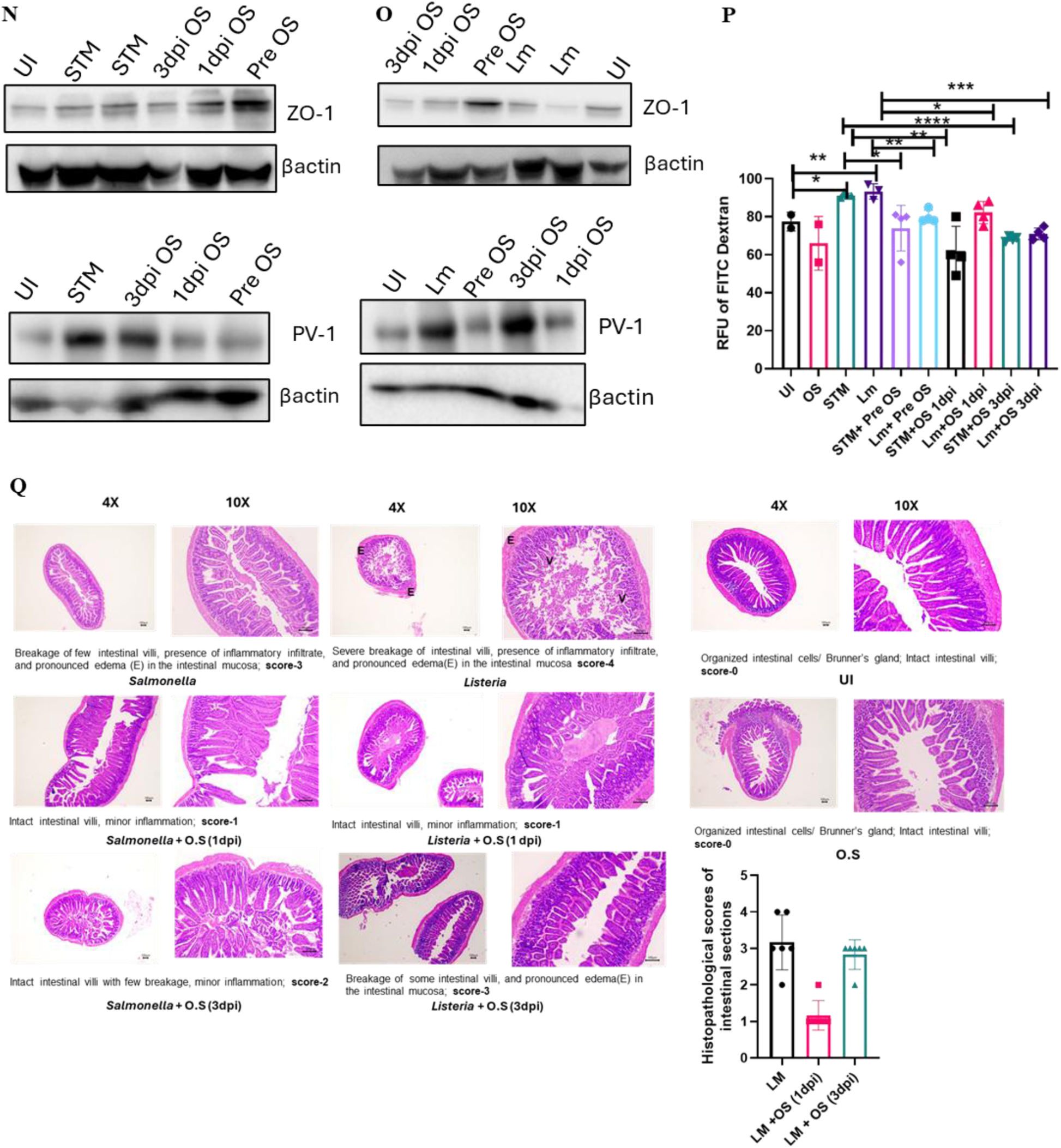
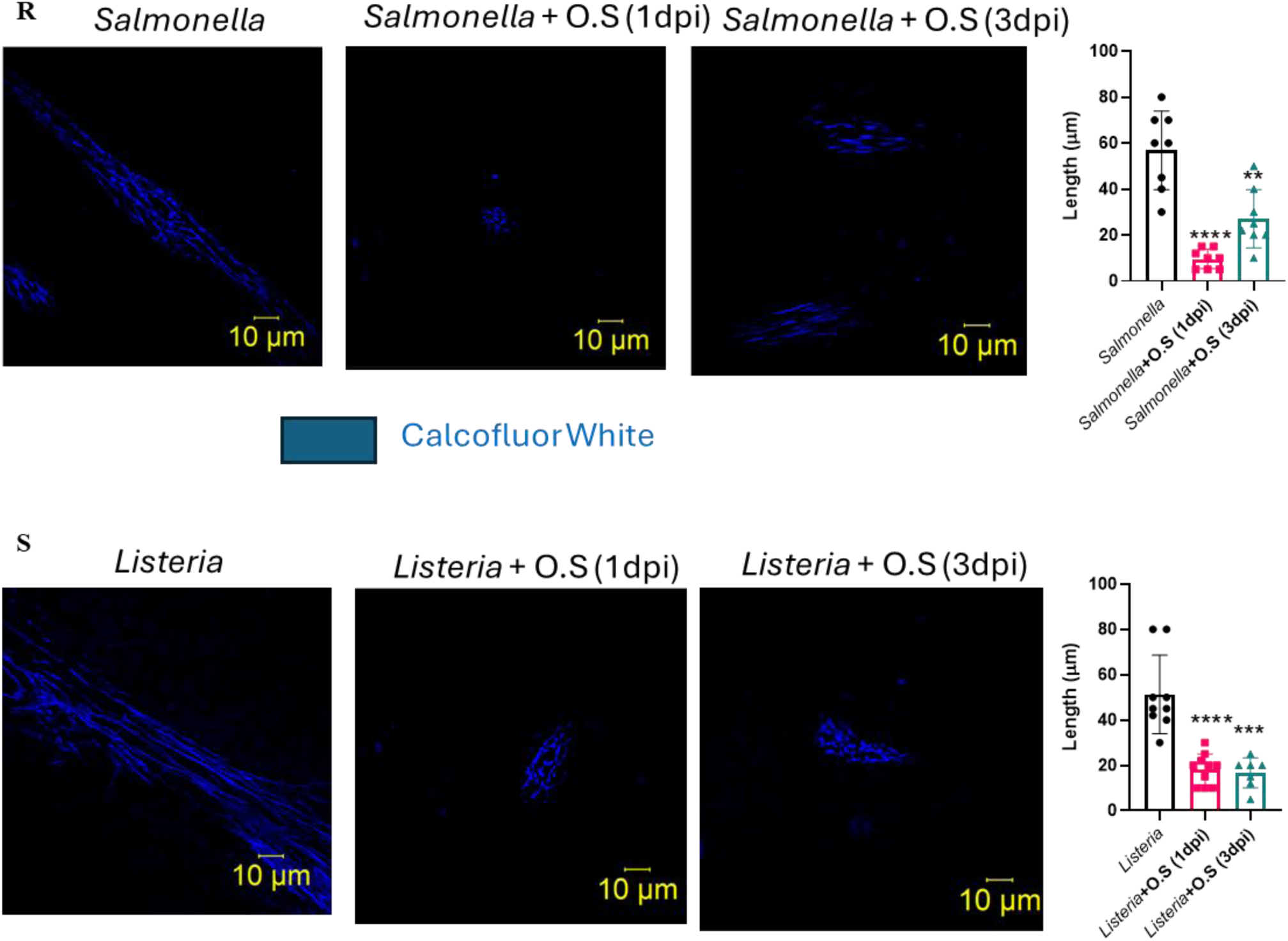
*Odoribacter splanchnicus* administration post-establishment of infection protected mice against *Listeria monocytogenes* and *Salmonella* Typhimurium. **A-C**- STM and LM organ loads in feces at 3^rd^ day post-infection (A), 5^th^ post-infection (B) and in ileal tissues (contents and wall) at 5^th^ day post-infection (C) upon OS administration at 1-dpi or 3-dpi. Data is representative of N=2, n=6 where N denotes number of biological replicates and n denotes the number of mice per group. Mann-Whitney Test was performed to obtain p-values. **D-H**-Quantitative gene expression data of several antimicrobial peptides (AMPs) precursor genes in LM and STM*-*infected mice intestinal tissues harvested at 5^th^ dpi upon OS colonization at 1-dpi or 3-dpi. Unpaired Student’s t-test is performed to obtain the p-values. Data is representative of N=2, n=4. **I-M**- Quantitative gene expression study of tight junction protein genes *Tjp1*, *Claudin5*, GVB damage marker *PV-1*, angiogenesis marker-*Vegf,* and *Axin2* in LM and STM-infected mice intestinal tissues harvested at 5^th^-dpi upon OS colonization at 1-dpi or 3-dpi. Unpaired Student’s t-test is performed to obtain the p-values. Data is representative of N=2, n=6. **N-O**- Immunoblotting representing the protein expression profile of ZO-1, and PV-1 in *Salmonella* (N) or *Listeria* (O)-infected mice intestinal tissue harvested on 5^th^-dpi with or without OS pre-or post-colonization (1-dpi or 3-dpi). Data is representative of N=3. **P**- Estimation of FITC fluorescence in mice plasma via gut permeability assay in *Salmonella* or *Listeria*-infected mice cohorts with or without OS pre-or post-colonization (1dpi or 3dpi) upon oral administration of FITC Dextran (25mg/kg) 4h before euthanization at 5^th^-dpi. Unpaired Student’s t-test is performed to obtain the p-values. Data is representative of N=2, n=4 (UI and OS, n=2). **Q-** Haematoxylin-eosin-stained intestinal sections harvested at 5^th^-dpi from Uninfected (UI) or *Salmonella* or *Listeria*-infected mice cohorts having non-administered or administered OS (at 1^st^ or 3^rd^ dpi). 0-represents normal pathology, 1- represents- mild pathology, 2-3-represents severe pathology. Data is representative of N=2, n=6. **R**- Representative Calcofluor white (CW) staining of STM*-*infected mice spleen tissues at 5^th-^day post-infection with or without OS administration at 1^st^ or 3^rd^ dpi depicting the quantification of the length of the fibril architecture of CW-stained sections. One-way ANOVA and multiple comparison test were used to obtain the p-values. **S**- Representative CW staining of LM-infected mice spleen tissues at 5^th-^dpi with or without OS administration at 1^st^ or 3^rd^ dpi depicting the quantification of the length of the fibril architecture of CW-stained sections. One-way ANOVA and multiple comparison test were used to obtain the p-values.

**Fig. S6.**
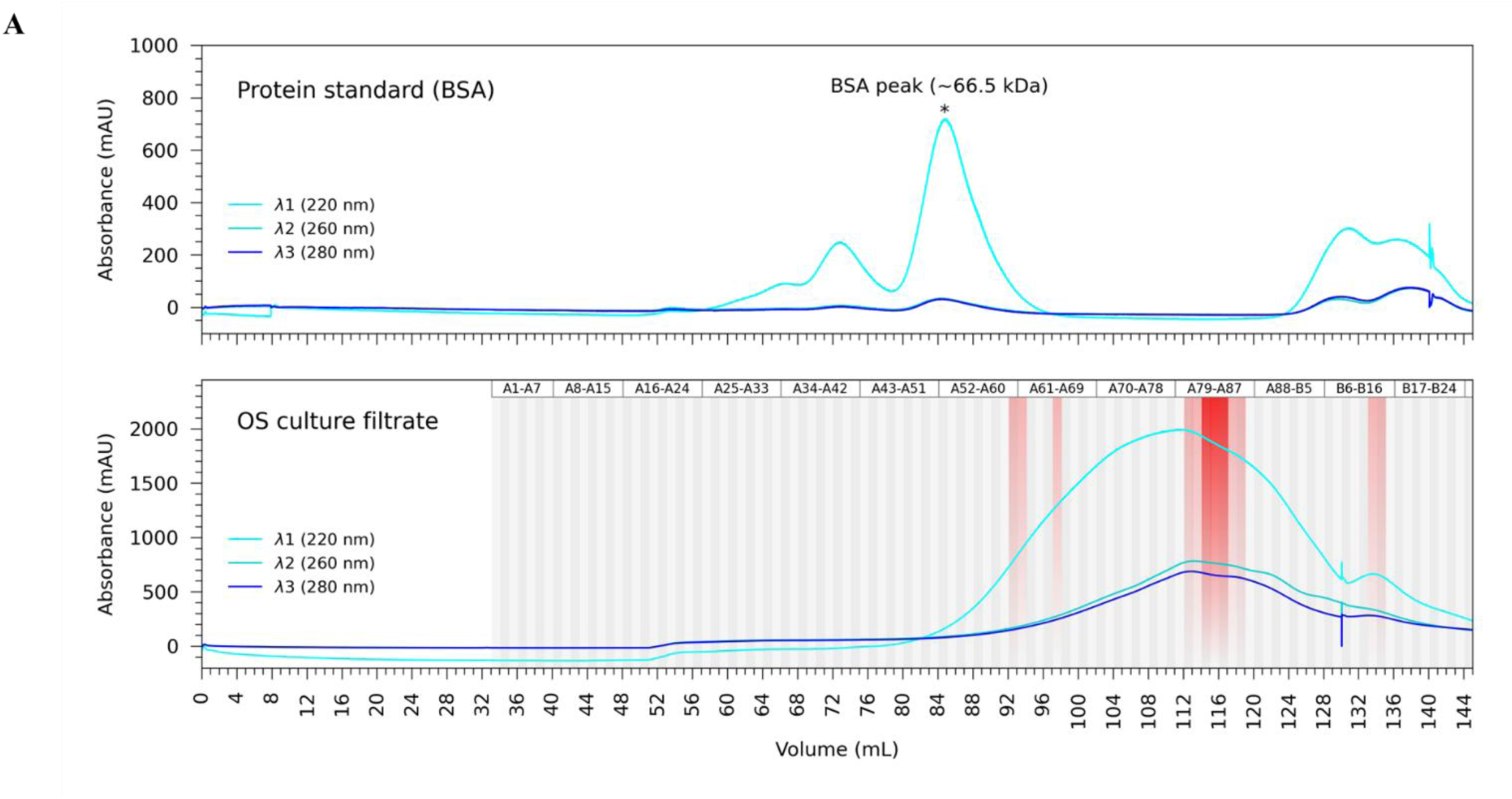
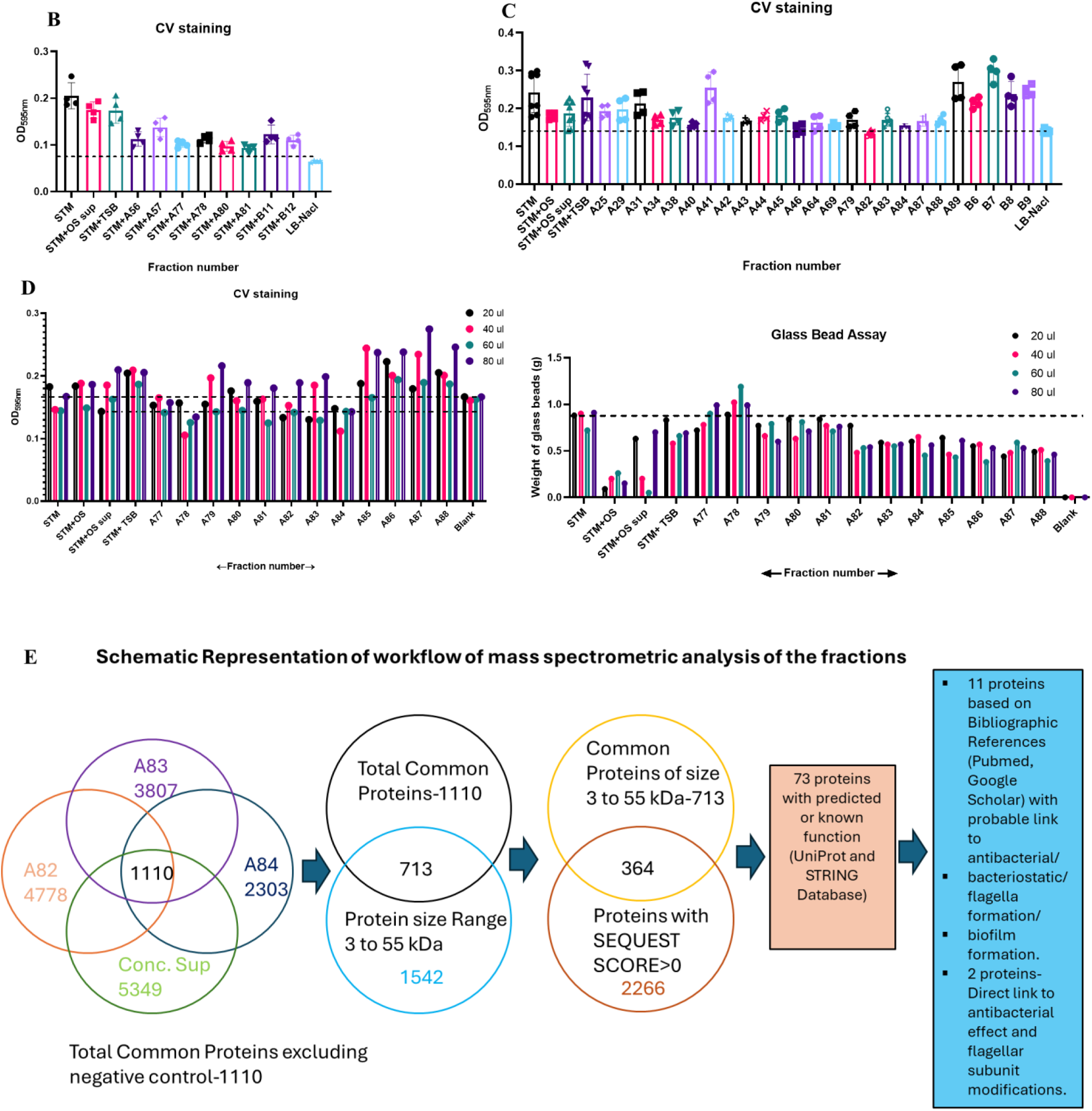
Characterization of the inhibitory active molecule within *O. splanchnicus* culture supernatant. **A-**Gel filtration chromatogram of protein standards and the OS culture supernatant. Red shaded region shows the fractions with biofilm inhibitory activity. The dark shaded region indicates the fractions with the highest inhibitory activity. **B-C**- Biofilm assay showing the biofilm formation ability of STM after 72h of inoculation in presence of OS supernatant gel filtration fractions. Data is representative of N=2, n=2. One-way ANOVA is used to obtain the p-values. **D**- Biofilm assay showing the biofilm formation ability of STM after 72h of inoculation in presence of varying concentration of OS supernatant biofilm inhibitory gel filtration fractions. Data is representative of N=2, n=1. **E**-Schematic representation of the workflow of mass spectrometric analysis of the gel filtration fractions (A82, A83, A84 and Concentrated supernatant).

**Fig. S7.**
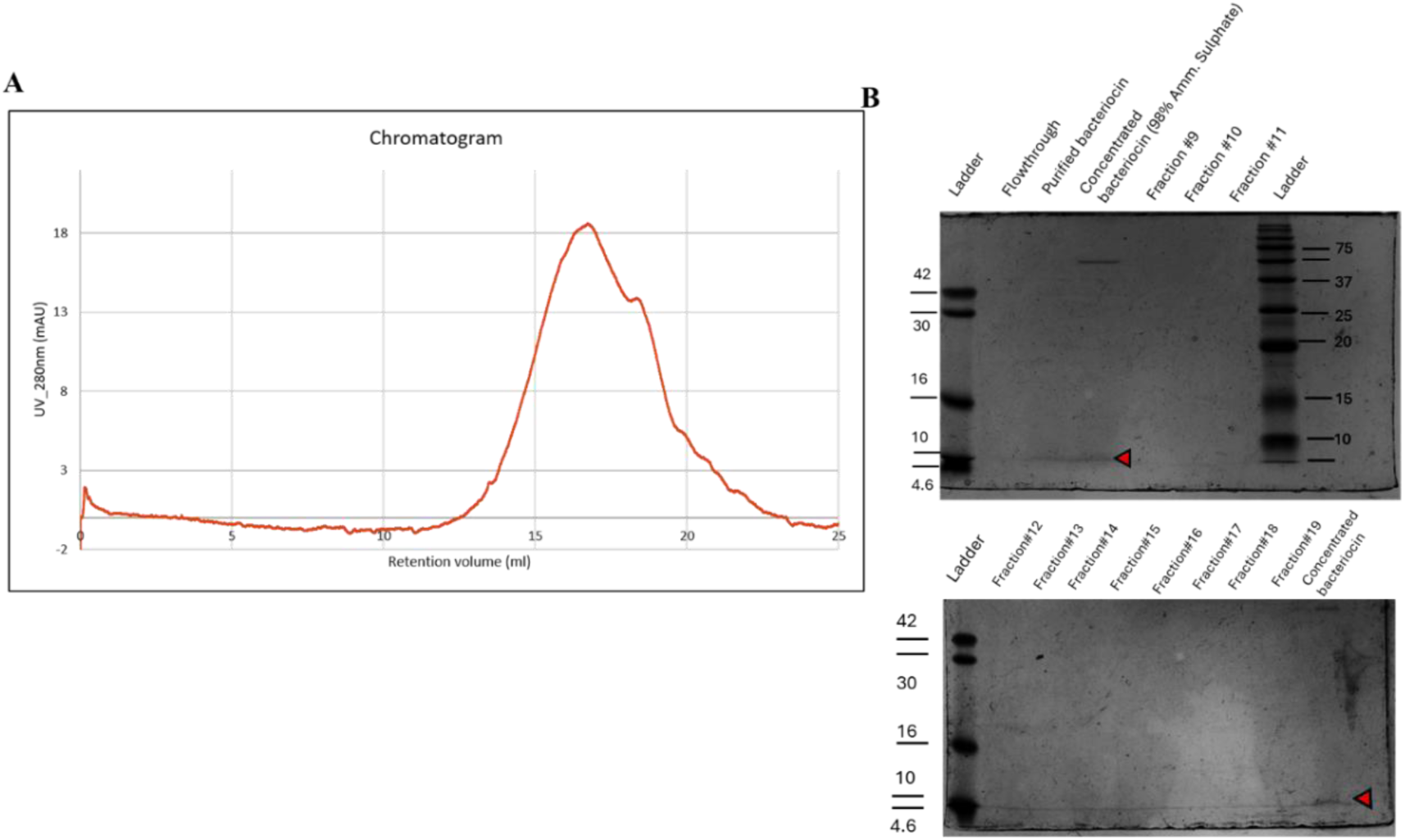
Purification procedure of *O. splanchnicus* produced bacteriocin via ammonium sulphate precipitation method followed by Fast Performance Liquid Chromatography. **A**-Size exclusion chromatogram of OS-produced bacteriocin fractions post 95-98% ammonium sulphate precipitation. **B**- Coomassie-stained SDS PAGE showing the purified and concentrated OS bacteriocin.

**Fig. S8.**
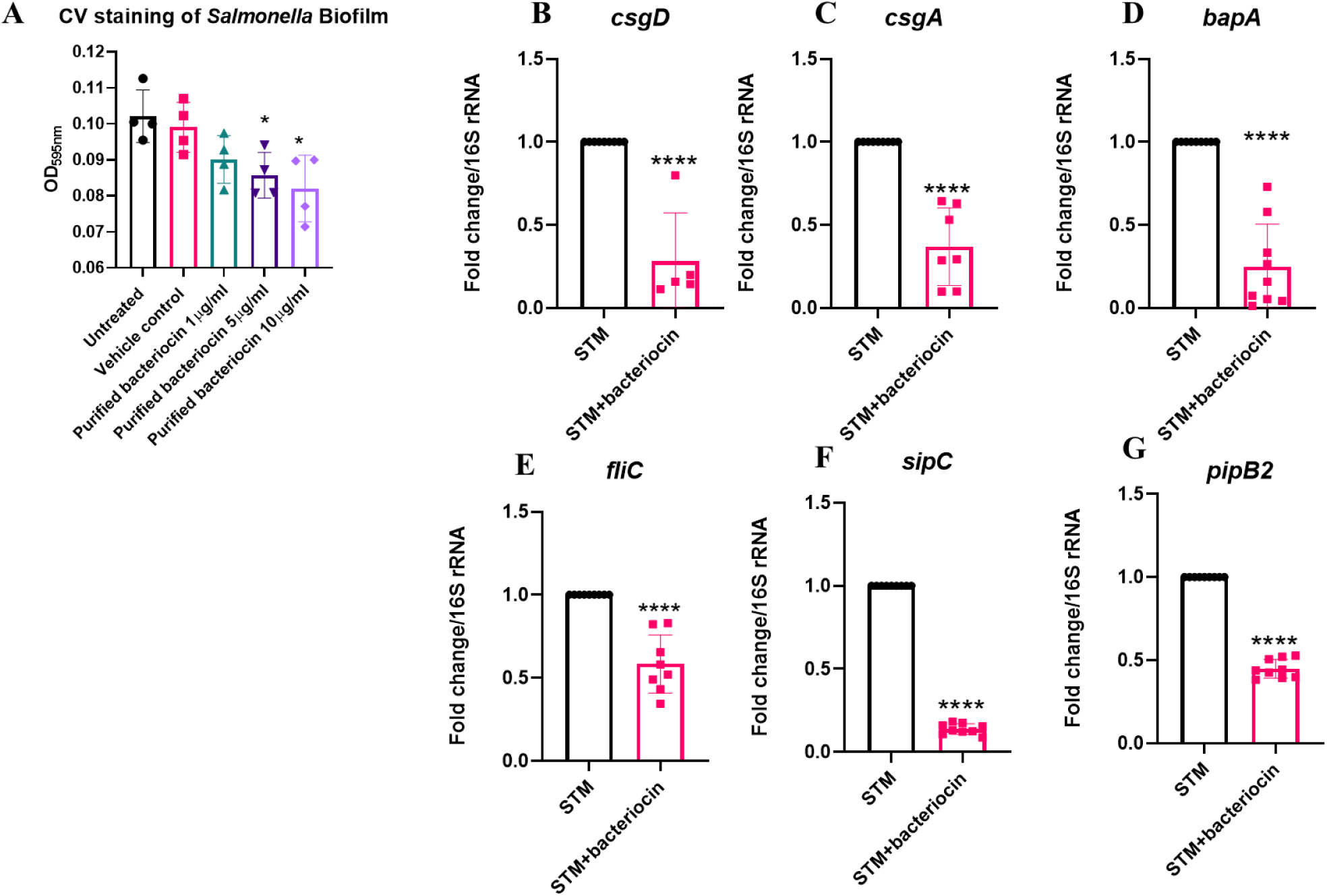
*O. splanchnicus* bacteriocin attenuates *Salmonella* pathogenesis *in vitro*. **A**- Biofilm assay showing the biofilm formation ability of STM after 72h of inoculation under varying concentration of OS bacteriocin. Data is representative of N=2, n=2. Unpaired t-test is used to obtain the p-values. **B-G**- RT-qPCR mediated gene expression of STM biofilm regulatory genes *csgD* (B), *csgA*(C), *bap* (D), along with flagellar gene *fliC*(E), SPI-1-*sipC* (F) and SPI-2-*pipB2*(G) in presence or absence of OS bacteriocin treatment. Data is representative of N=2, n≥5. Unpaired Student’s t-test was used to obtain the p-values.

**Table S1.** Selected List of 11 proteins from Mass Spectrometric Analysis based on bibliographic search.

| Accession ID | Protein Name and | Referenced Function |
| --- | --- | --- |
| A0A412WBT8 | Immunity protein Imm5 domain-containing protein | Uncharacterized |
| F9Z8G5 | Peptidase | Modulate biofilm formation and host defense. |
| R6F650 | Carboxyl-terminal protease | Critical for post-translational processing and associated with regulation of gene expression, stress response, maintenance of cell envelope integrity and bacterial virulence. |
| A0A412TJE3 | ATP-dependent Clp protease proteolytic subunit | Influence virulence and infectivity of pathogenic bacteria. ClpP enhance biofilm formation in *Enterococcus faecalis*. |
| F9Z604 | Peptidase C39 bacteriocin processing | The cysteine peptidase cleave the double glycine leader peptides from the precursors of bacteriocin |
| A0A412TJV3 | UDP-2,4-diacetamido-2,4, 6-trideoxy-beta-L-altropyranose hydrolase | It catalyzes the third step in the biosynthesis of pseudaminic acid. Pseudaminic acid is a sialic-like sugar that modifies flagellin. |
| A0A412WSU3 | Bacteriocin | Antimicrobial peptides that can kill or inhibit closely related or unrelated bacterial strains. |
| A0A413I832 | Serine Protease | Extracellular proteases regulate biofilm formation and growth of competitors. |
| A0A412W3J2 | Glutamate dehydrogenase | Involved in bacterial response to stress adaptation and virulence. |
| A0A412WSS0 | Bacteriocin | Antimicrobial peptides that can kill or inhibit closely related or unrelated bacterial strains. |
| F9ZBU9 | Peptidylprolyl isomerase | Promote biofilm formation in *Mycobacterium tuberculosis*. |

**Table S2.**
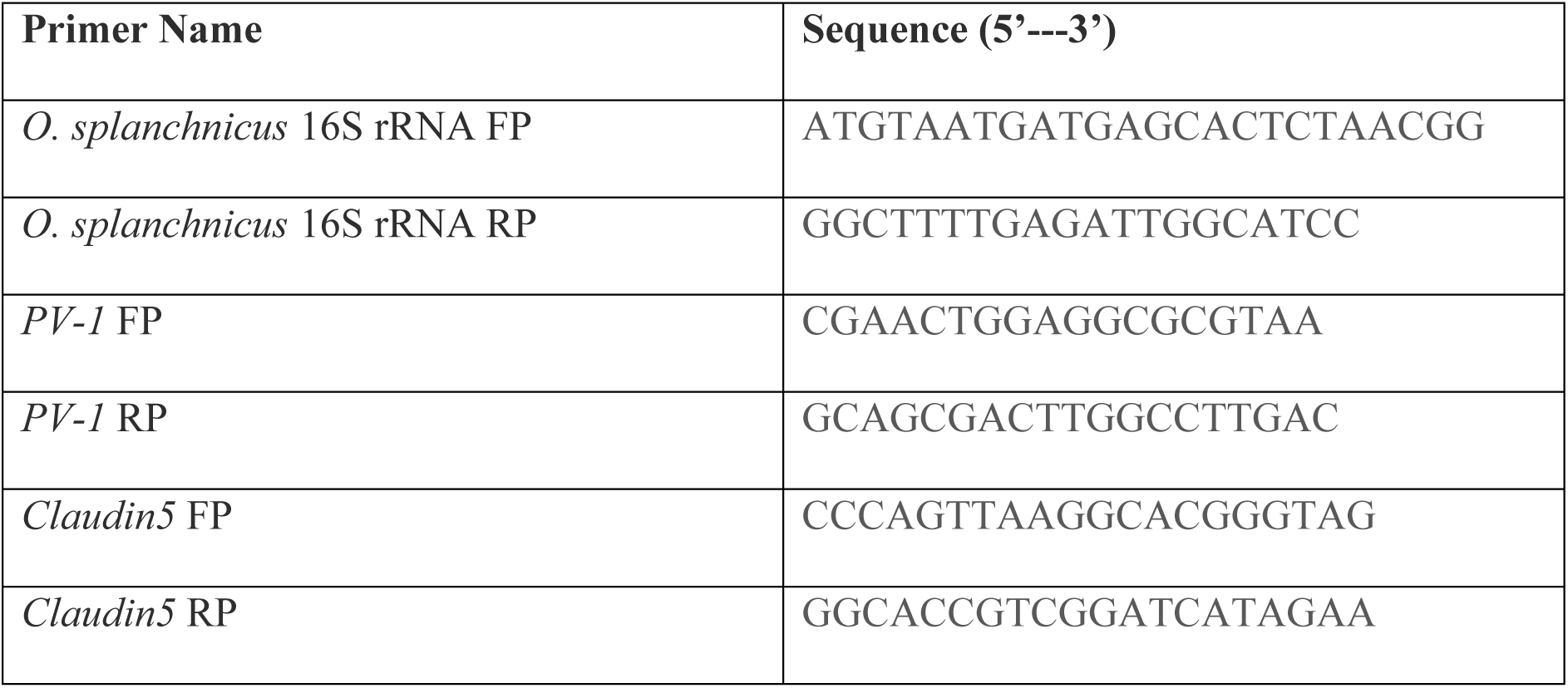
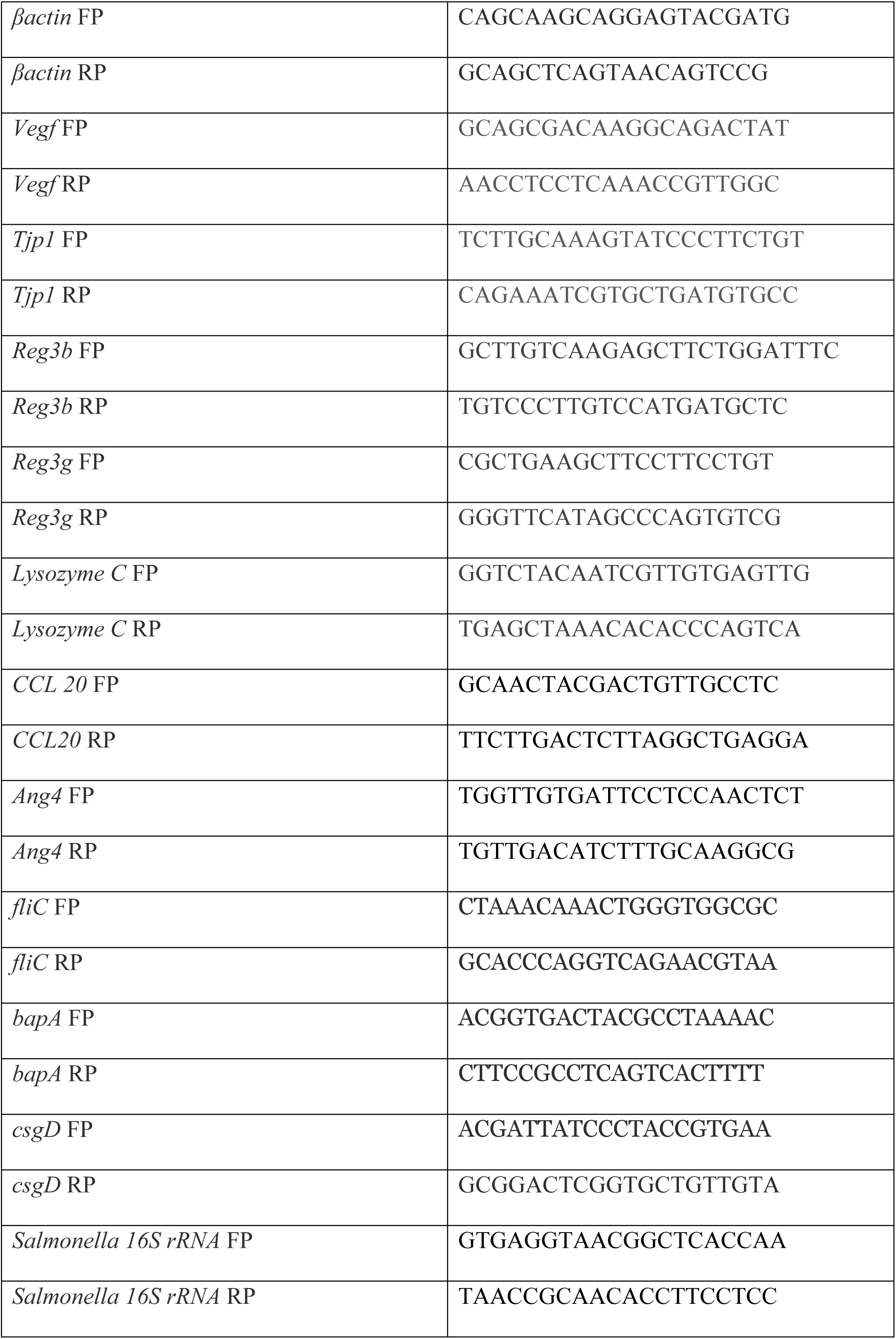
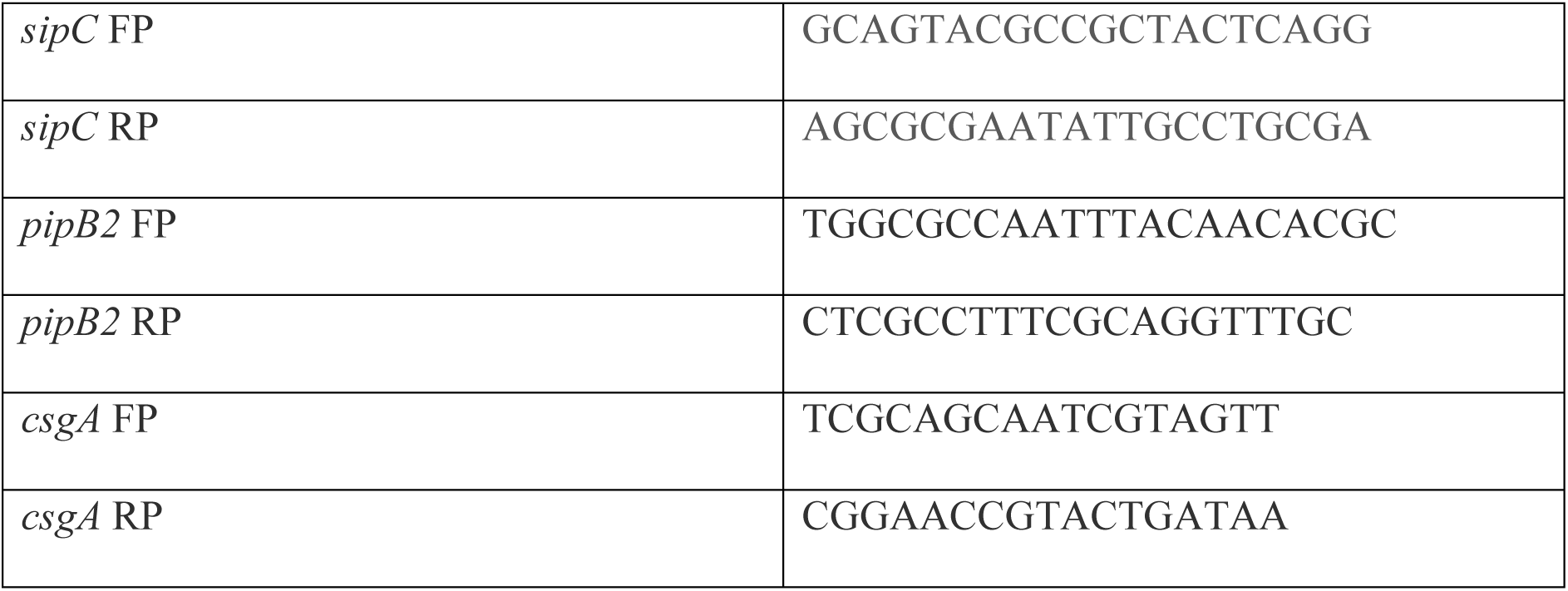
List of primers.

**Table S3.** List of Antibodies.

| **Antibody Name** | **Catalogue Number** | **Source** |
| --- | --- | --- |
| PV-1 | **A15906-20ul (Abclonal)** | **Rabbit** |
| ZO-1 | **R26.4C DHSB, CRL 1597** | **Mouse** |
| $\beta$ actin | **A3854 (Sigma), Clone AC-15** | **Mouse** |

